# A few genes modulate the ecological impact of a freshwater fish species on food webs and ecosystem functions

**DOI:** 10.64898/2026.10.20.760412

**Authors:** Maxime Stanislawek, Patrik Nosil, Allan Raffard, Laura Fargeot, Murielle Richard, Madoka Krick, Camille Poesy, Maxim Lefort, Géraldine Loot, Simon Blanchet

## Abstract

Genetic diversity within species can shape food webs and ecosystem functioning as strongly as species diversity, yet the genomic architecture underlying these effects remains unclear. Gene-to-ecosystem relationships may arise from genome-wide diversity, from a limited number of ecologically-important genes, or a combination of both. Experimental tests of these hypotheses are lacking. Here, we manipulated genomic diversity of a top-predatory fish in mesocosms under ambient or warm conditions, and quantified ecological responses spanning fish biomass, prey communities and ecosystem functions. Genome-wide diversity explained few ecological responses, with moderate effects (*r* = 0.260). In contrast, variation at specific candidate SNPs predicted all ecological responses consistently across climatic conditions, with effect sizes being more than twice those of genome-wide estimates (*r* = 0.599 ). Moreover, combining information from small subsets of candidate SNPs outperformed both single-locus and genome-wide diversity for all ecological responses (*r* = 0.841), consistent with an oligogenic architecture for gene-to-ecosystem relationships. Enrichment analyses revealed functional pathways related to growth, development and metabolic regulation, driven by a small set of pleiotropic genes. We demonstrate that the ecological consequences of intraspecific genetic variation arose from a limited fraction of the genome, highlighting the importance of functional genomic structure, rather than genome-wide diversity alone, in shaping eco-evolutionary dynamics.

## INTRODUCTION

Understanding the relationships between biodiversity, food web dynamics and ecological processes is critical for predicting the ecosystem-scale consequences of ongoing changes in biodiversity. Species diversity strongly influences food web dynamics, as well as key ecological functions such as productivity or nutrient retention, and promotes their stability across time and space^1–3^. Moreover, there is growing evidence, -from diverse and independent research fields, that diversity measured within a single species (intraspecific diversity) can alter food webs and support ecosystem functions as much as species diversity^4–6^. For instance, different morphotypes of a predatory species can have substantial diverging effects on their prey communities^7^ -see also^8^ for a herbivore-plant system study-, whereas populations with greater genetic diversity generally display higher and more stable biomass than genetically depleted populations (a pattern akin to biodiversity-function relationships^9,10^). Altogether, these findings imply that species loss may not be a prerequisite for ecosystem-level impacts; even cryptic genetic or phenotypic changes within a single species can substantially affect food webs and the whole ecosystem functioning. Given the observed and predicted loss in genetic diversity worldwide^11,12^, it becomes urgent to better grab the relationships between genetic diversity and ecosystem functioning (i.e., “gene-to-ecosystem relationships”).

Despite evidence of the ecological impacts of intraspecific genetic diversity across a wide range of taxonomic and ecosystem contexts^4,5,13^, the functional bases underlying gene-to-ecosystem relationships remain poorly understood. Two genetic-based hypotheses have been proposed to explain gene-to-ecosystem relationships, yet it remains unclear which one prevails. On the one hand, population genomic diversity measured at the genome-wide scale has been assumed to explain a large proportion of the trait diversity involved in ecological processes^14^. Under such a hypothesis (hereafter termed the “genome-wide hypothesis”), genetically diverse populations may display higher functional and trophic complementarity among individuals, leading to more efficient resource partitioning, resource use and hence energy transfer across trophic levels^14,15^. This hypothesis has been tested across a wide range of taxa, from plants to animals, with consistent positive effects of genome-wide diversity on biomass productivity, resilience and stability^6,9,10,16,17^. On the other hand, it has recently been proposed that variation at a few “ecologically important genes” (or “keystone genes” when these genes have disproportionally large effects) may sustain such ecological effects of intraspecific genetic diversity^18–20^. This hypothesis (hereafter the “local genomic hypothesis”) emerged from a series of founder studies on *Populus* genotypes^13,21,22^, stipulating that local genomic effects could result in competitive functional traits with disproportionate consequences for food webs and ecosystem functions. One convincing example of a local genomic effect is that of Barbour et al. (2022) who showed in *Arabidopsis thaliana* that allele frequency at a single growth-related gene (*AOP2*)^19^ indirectly promoted species coexistence in a three-species food web ^see^ ^also, 23,24^. However, although a powerful case study, this study was restrained to a highly controlled and simplified food-web system, so it remains unclear to what extend a few genes can actually be important for ecological dynamics under realistic conditions. Moreover, very few studies have directly compared the likelihood of these genome-wide versus local genomic hypotheses concerning gene-to-ecosystem relationships, a necessary step for setting realistic models coupling ecological and evolutionary dynamics.

These two key hypotheses rely on gene-to-phenotype linkages, which are often environment-dependent, allowing identical gene pools to produce different phenotypes under varying environmental conditions. Therefore, the ecological impacts of intraspecific diversity are also expected to depend on environmental contexts^14,23^. Moreover, stressful environments may amplify the influence of genetic diversity, by revealing “cryptic” variation that remains hidden under baseline conditions, both at the local and genome-wide scales^14,25^. For instance, Prunier et al. (2023) showed that stressful eutrophic conditions triggered the positive effect of the genome-wide diversity of a single fish species on the temporal stability of fish biomass over time^10^. Yet, despite these findings, our understanding of how genetic diversity interacts with environmental stressors, such as climate change, to influence ecosystem functioning remains limited. This is concerning, given that biodiversity loss and climate change are two main components of the ongoing ecological crisis.

Here, we tested whether genome-wide diversity (genomic diversity across all measured loci at the genome scale) or local genomic diversity (genomic diversity measured at the scale of single or subsets of genomic loci) better explains the effects of intraspecific diversity on ecological dynamics under climate change. The occurrence of strong local genomic effects would suggest the existence of ecologically-important genes that could be used to predict the ecological outcomes of genetic diversity loss. To do so, we conducted a realistic year-long mesocosm experiment in which we manipulated the genetic diversity of a top-predatory fish in different temperature treatments. In the experiment, we monitored fish populations, their prey communities, and ecosystem functions to assess the effects of genetic diversity across multiple biological levels and entire food webs under climate change scenarios. Specifically, we compared the relationships between genomic diversity measured at the genome-wide scale *versus* at the single-locus scale and ecological responses. We expected associations at both scales, but predicted that local effects would on average be stronger than genome-wide effects, consistent with the existence of ecologically-important genes^18,19,24^. We further hypothesized that the strength of genomic diversity effects would be context-dependent, with stronger effects under warming stress^10^. Finally, we investigated the functional roles of loci linked to ecological responses to gain a mechanistic understanding of how intraspecific diversity shapes ecosystem dynamics.

## MATERIALS AND METHODS

### Biological model and experimental design

The minnow (*Phoxinus dragarum*) is a small Cyprinid fish species measuring 4 to 8 cm in length as an adult. This is an abundant species living in mid-altitude streams and rivers in South-Western France, as well as in Pyrenean Mountain lakes where it has been introduced. It is a generalist species feeding on small invertebrates, algae and zooplankton, and can have strong top-down effects on lower trophic levels and ecosystem functions^26,27^

The experimental design involved manipulating the climate of experimental ponds and the genetic diversity of *P. dragarum* populations. The design has been previously described in Stanislawek et al. (2025) to which readers can refer for further details^28^. In Stanislawek et al. (2025), we tested the relative influence of climate warming and fish genetic diversity (measured only at the genome-wide level) on the size structure of experimental fish populations^28^. Here, we build on the same experiment, but we investigate the genomic underpinning of the ecosystem impacts of *P. dragarum* by considering the entire food web, from the fish to the primary producers. Briefly, we used 24 circular tanks (∼2000 L in volume) from the Aquatic Metatron^29^ filled with filtered water and in which we added a 3 cm-layer of small pebbles. Six months before the start of the experiment (May 2019) tanks were inoculated with zooplankton (copepods, cladocerans, rotifers), phytoplankton (primary producers), molluscs (*Lymnaea fluviatilis*, n=20 per tank), amphipods (*Gammarus ariegiensis*, n=50 per tank) and isopods (*Asellus aquaticus*, n=50 per tank). Shelters (bricks and coconut brushes, 2 per tank) were added and the tanks were kept open for natural colonization by winged invertebrates (mayflies, dipteras, beetles…).

From October 2019, 12 out of the 24 mesocosms were artificially heated by +3°C above ambient temperature using a controlled supervision system^29^ and leading to two climatic treatments: ambient vs. warm temperature. The genetic diversity of experimental fish populations was manipulated by introducing different combinations of individuals from genetically divergent wild populations^28^. The wild populations used as “source” populations were captured by electrofishing in summer 2019 from 6 rivers in southwestern France (Lez, Hers, Gouarège, Salat, Arize and Volp Rivers, see Table 1 in^28^ for exact geographic coordinates and further details on each population). We sampled both adults and juveniles (∼50 specimens each) from each river that we maintained separately for ∼3 months in 1000L tanks before the start of the experiment. In October 2019, individuals were introduced from either two source populations (which is expected to result in populations with “low” genetic diversity) or four source populations (which is expected to result in populations with “high” genetic diversity) out of the six source populations. We varied the composition of each experimental population (resulting in 12 combination) and each combination was subjected to the two climatic treatments. Each experimental population consisted of 8 juveniles and 12 adults that were anesthetized and weighed to the nearest 0.01 g before introduction into the mesocosms. A pelvic fin sample was also collected from each individual (n = 480) for genomic analyses. We did not notice birth during the experiment. The experiment was run according to ethical rules under agreement number APAFIS #34179-2021120109429216 v5.

**Table 1.**
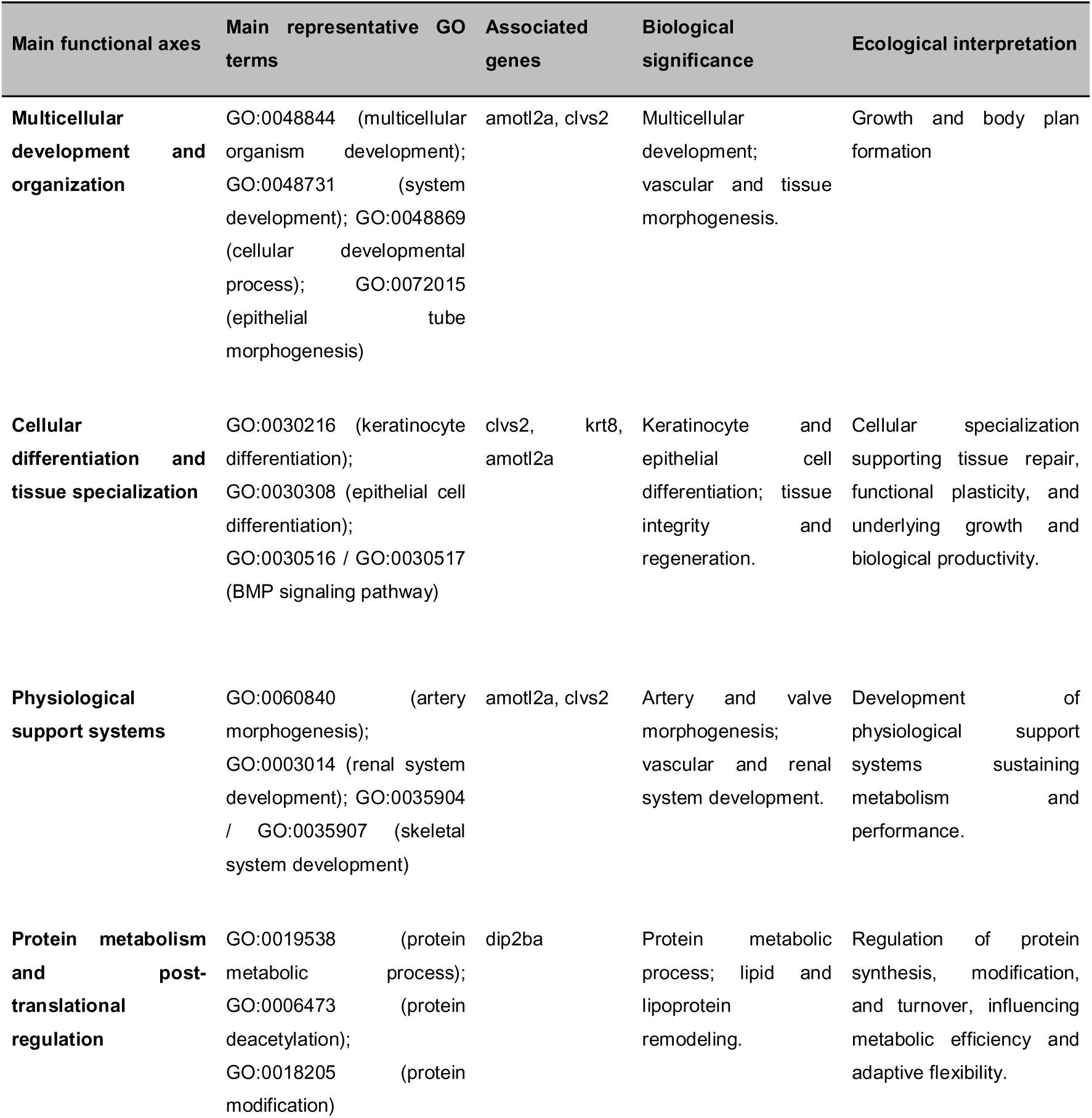
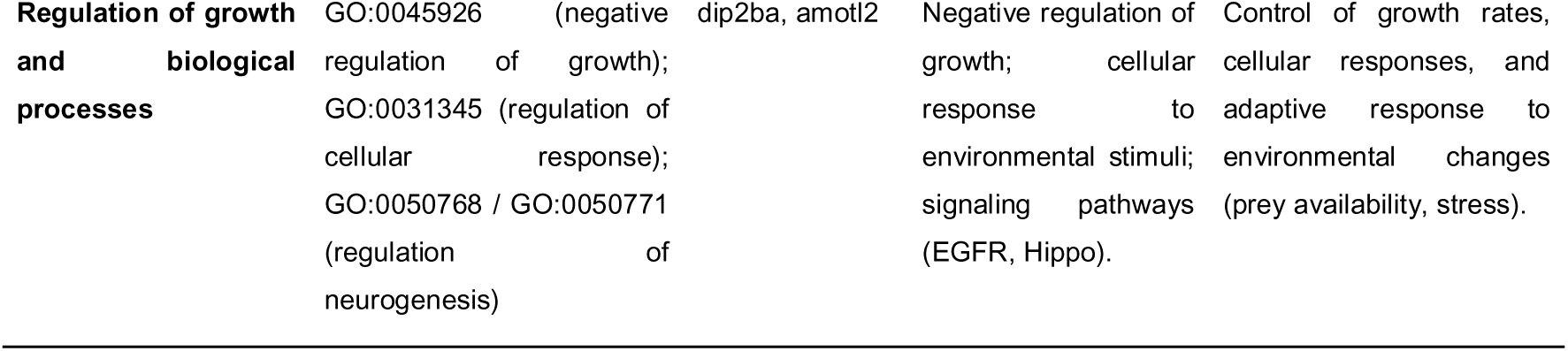
Main functional axes associated with candidate SNPs underlying ecosystem effects. Gene Ontology (GO) enrichment analysis performed on genes associated with candidate SNPs identified five major functional axes related to organism development, tissue differentiation, physiological support systems, protein metabolism, and growth regulation. Representative enriched GO terms were grouped into biologically coherent functional axes. Associated genes correspond to proteins annotated from candidate SNPs located within coding sequences or promoter regions. Biological interpretations summarize the main functions represented by the enriched GO terms, whereas ecological interpretations propose the potential mechanisms through which genomic variation may influence organismal performance and, ultimately, ecosystem processes.

#### Genomic data

We assessed the genomic diversity of each individual using bi-allelic Single Nucleotide Polymorphism (SNP) markers gathered at the genome level (see^28^ for details). The DNA was extracted from each fin (Blood & Tissues kits, Qiagen®), and was sent to Diversity Arrays Technologies Ltd, in Australia (www.diversityarrays.com) for the genotyping of SNPs using a reduced-representation sequencing approach (DArT sequencing^30^). SNP calling was done by Diversity Arrays Technologies Ltd. based on a *P. dragarum* reference genome^10^. Markers with inappropriate quality control parameters, including a call rate ≤ 85% and missing data ≥ 20%, were removed. After filtering, 15799 SNP markers were retained for subsequent analyses. From that set of markers, we estimated for each experimental population the proportion of polymorphic SNPs, which we use as an estimate of *genome-wide* genomic diversity. As expected, experimental populations seeded from two source populations had a lower proportion of polymorphic SNPs compared to experimental populations seeded from four source populations (Welch t-test = 4.0176, df = 12.887, p = 0.001), with an overall gradient of global genomic diversity created across populations (see Figure 2 in^28^). Note that we also tested the association between genome-wide diversity and each ecological response using alternative estimates of diversity (the expected heterozygosity and a genetic distance estimating the number of SNPs by which two individuals differ), and results were similar to those found with the proportion of polymorphic SNPs (see the Results and Table S1). We conserved the percentage of polymorphic SNPs because overall fits were slightly better. For each SNP and each experimental population, we also estimated the allelic frequency of one of the two alleles in the experimental population as an estimate of the local genomic diversity. For a given SNP locus, allele frequencies close to 0.5 indicate experimental populations that are more diverse than populations displaying allele frequencies close either from 0 or 1 (indicating monomorphic populations).

**Figure 1.**
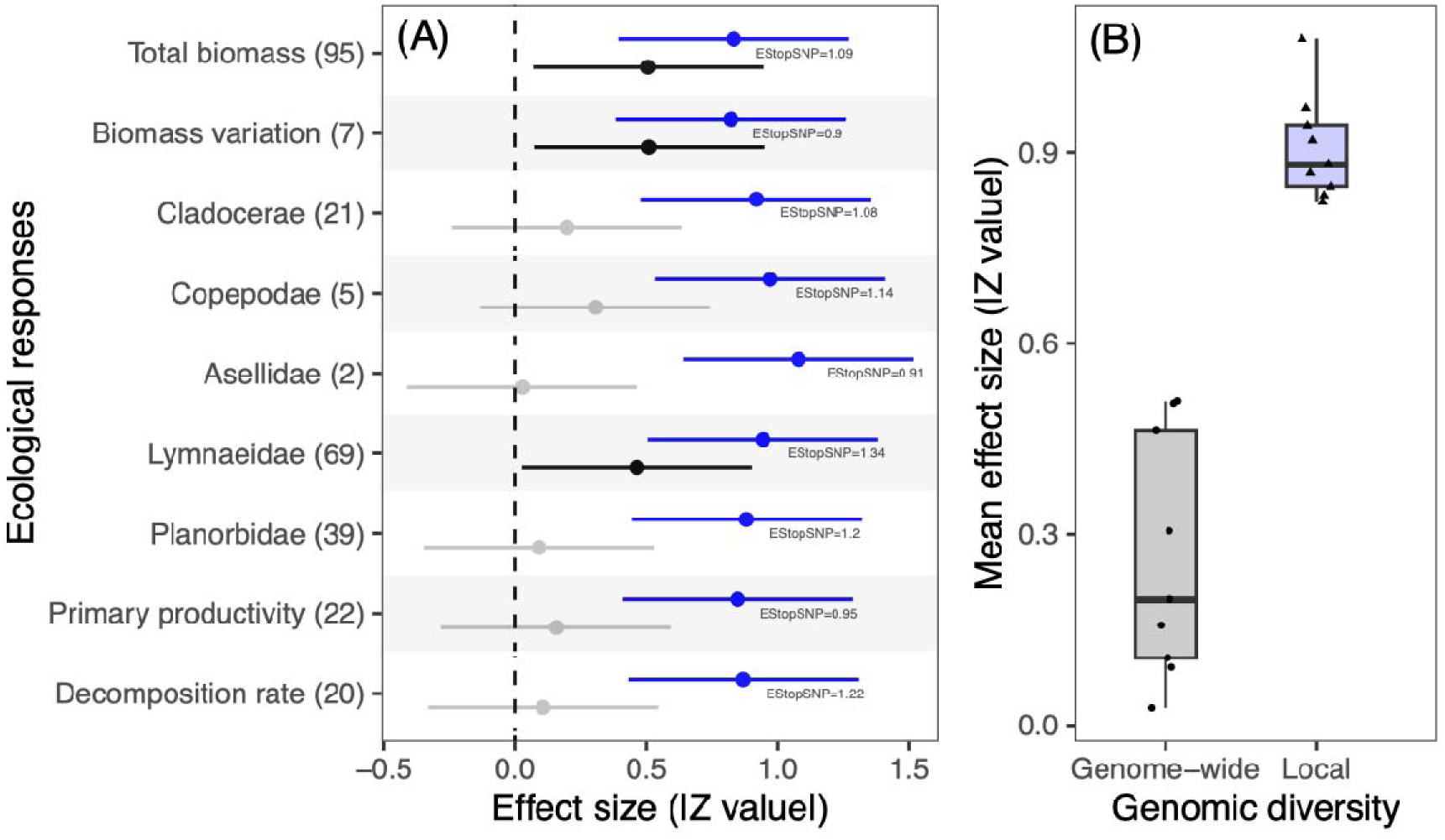
Comparison between local genomic effect sizes and genome-wide effects. (A) Absolute effect sizes (Z-values with 95% CI) for genome-wide effects (grey and black dots, black dots indicating significant genome-wide effect sizes) and local genomic effects (i.e., effect sizes averaged across candidate SNPs, blue dots). Effect sizes are shown for each of the nine ecological responses, with the number of candidate SNPs (FDR<0.05) indicated for each of them. “ES_topSNP_” stands for the effect size of the top candidate SNP for each ecological response, i.e., the SNP with the highest effect size among the set of candidate SNPs. (B) Boxplots contrasting the distribution of effect sizes for genome-wide (grey boxplot) and local genomic (blue boxplot) effects.

**Figure 2.**
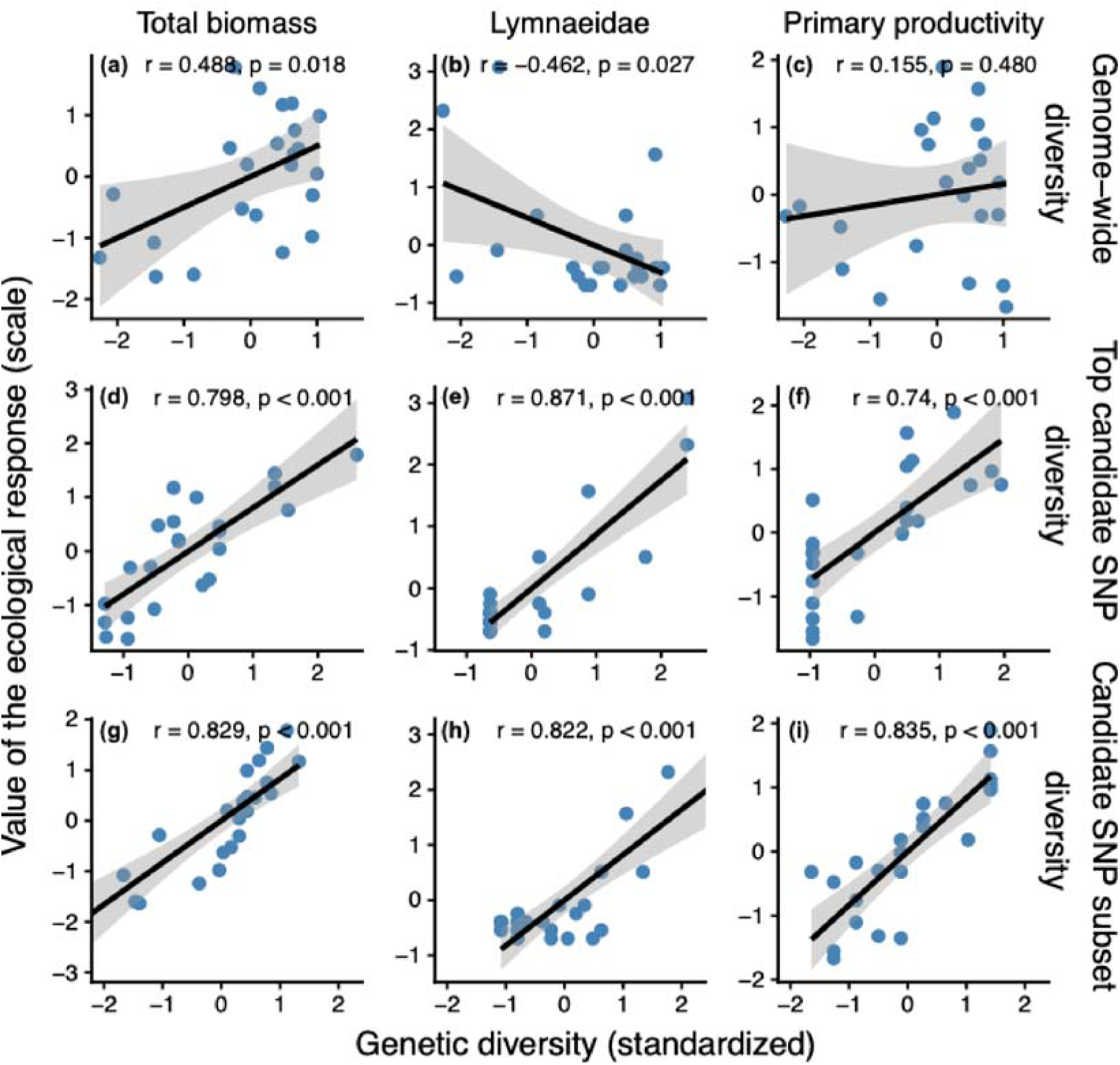
Relationships between genetic diversity in experimental population of *P. dragarum* and three out of the nine ecological responses (fish total biomass, prey Lymnaeidae abundance, estimate of phytoplankton primary productivity). Relationships are shown for different estimates of genomic diversity (raws): the genome-wide diversity (a-c, « Genome-wide diversity », expressed as the proportion of polymorphic SNPs across 15799 SNPs), the top candidate SNP genomic diversity (d-f, « Top candidate SNP diversity », expressed as the proportion of the major allele in each minnow population), and the candidate SNPs subset diversity estimated from an optimized subset of candidate SNPs (g-i, « Candidate SNP subset diversity », expressed as the proportion of polymorphic SNPs across the subset of candidate SNPs. In all biplots, estimates of genomic diversity have been scaled to the mean to make easier the comparison, and the highest positive values indicate the more diverse minnow populations. In particular, for the local effects of the top candidate SNPs, positive values indicate populations with an even allelic distribution whereas extreme negative values indicate monomorphic populations.

### Experimental survey and responses variables

We estimated ecological responses at three biological levels spanning the entire food web: the fish population, the prey community and the ecosystem functioning.

#### Fish population level

We used the fish biomass and its coefficient of variation within each tank respectively as estimates of fish productivity and population size structure (populations with greater genomic diversity should have higher biomass and CV as they should be more diverse phenotypically and exploit available resources more efficiently). Individuals were weighed three times to provide average estimates over the whole experiment: the initial measurement (T0), a second measurement after 7 months of experience (T1, May 2020), and a third measurement after 12 months of experience (T2, October 2020). Surviving fish were recaptured using baited minnow traps placed in each of the tanks until no fish entered the traps and no fish were observed swimming into the tank (recapture rate=98%). Survival rate averaged ∼70% at T1 and ∼40% at T2^28^. Captured individuals were anesthetized and weighed to the nearest 0.01g before being released back to their tank. For each tank, the fish biomass (sum of the biomass of all captured fish) was estimated at each time step and then averaged across the three-time steps to get a single integrative estimate. In the same way, the coefficient of variation of individual body mass was estimated at each time step and averaged across the three-time steps.

#### Prey community level

We focused on two categories of prey commonly consumed by minnows^26,27^: pelagic zooplankton and benthic macroinvertebrates. For each group, we estimated the abundance of preferred prey taxa within each tank, which can be assimilated to an estimate of secondary productivity. To identify the prey taxa targeted by minnows, we compared the abundance of each potential prey group between tanks containing fish and eight control tanks without fish (although not described above, these control tanks were set according to the same protocol and at the same time). Taxa showing significantly lower abundances in fish tanks relative to control tanks were considered as preferential prey. Based on this criterion, the main prey groups of zooplankton were Copepoda and Cladocera, while the main benthic prey groups were Asellidae, Lymnaeidae, and Planorbidae, which is coherent with the minnow biology. For these taxa, abundances were typically 5- to 10-fold lower in tanks with fish compared to control tanks.

Zooplankton abundance was quantified by filtering 5 L of water through a 200-μm mesh sieve at both T1 and T2. Samples were preserved in 70% ethanol. Adult Copepoda and Cladocera individuals were subsequently counted under a microscope, and counts were averaged across the two sampling times. The total number of individuals per group and per tank was used as a proxy for cladoceran and copepod productivity.

Benthic macroinvertebrates were sampled at T2 only. Organisms were collected from a standardized substrate area (50 cm²) using a hand net (mesh size: 0.5 mm), as well as from artificial shelters (bricks and coconut brushes) placed in each tank. Samples were preserved in 70% ethanol, then sorted and identified to the family level before counting. The total number of individuals per targeted taxon and per tank was used as an estimate of the productivity of Asellidae, Lymnaeidae, and Planorbidae.

#### Ecosystem functioning level

We focused on two key ecosystem functions: the primary productivity of pelagic microalgae and the decomposition rate of detrital organic matter. These two functions are partly regulated by the diversity and abundance of prey consumed by minnows, and we therefore expected them to vary indirectly with the genomic diversity of fish populations^31^.

Primary productivity was estimated using chlorophyll-a concentration (mg·L⁻¹) in the water column, measured weekly with a fluorometric probe (AlgaeTorch, bbe, Germany). Measurements (n = 52 per tank) were averaged over the duration of the experiment to provide an integrative estimate of pelagic primary productivity.

Decomposition rate was assessed at T1 and T2, as the mass loss of standardized leaf litter from *Alnus glutinosa*, a common riparian tree species^6^. At each time point, two coarse-mesh litter bags (1 × 1 cm mesh), each containing 4 g of air-dried *A. glutinosa* leaves, were placed in each tank. After approximately 40 days, litter bags were retrieved, and the remaining leaf material was oven-dried (60°C for 48 h) and weighed to the nearest 0.001 g using a precision balance (Ohaus®). Decomposition rate was calculated as the daily loss of leaf mass. Values were averaged over the two replicated bags and the two sampling periods (T1 and T2).

## Statistical analyses

One mesocosm in which an abnormally high fish mortality rate was observed was excluded from the dataset prior to subsequent analyses (N_final_=23 replicates). Specifically, the proportion of dead individuals after 12 months varied from ∼30 to 60% across all other tanks, whereas in the excluded tank, all fish died suddenly after month 10. All statistical analyses were performed using the R software (version 4.2.3).

### Comparing genome-wide and local genomic effects

We tested the effect of genetic diversity at both the genome-wide scale (i.e., the proportion of polymorphic SNPs) and the local scale (i.e., the allelic frequencies of each SNP independently) for each ecological response separately. To do so, we used generalized linear mixed models (GLMMs, glmmPQL function, MASS R package) to estimate the strength (effect size, see hereafter) and the significance of the relationships between genetic diversity (at the genome-wide or local scale) and each ecological response. We included the composition of experimental populations (i.e., the populations used to source each tank) within tanks as a random effect in the models to control for identity effects. We assumed a QuasiPoisson error-term distribution to fit abundance (count) data and to account for possible excesses of zero (Copepods, Cladocerans, Asellidae, Lymnaeidae and Planorbidae) and a Gaussian error-term distribution for other ecological responses (total biomass, CV of individual biomass, decomposition rate and primary productivity).

To assess the effects of genome-wide diversity, we related the proportion of polymorphic SNPs within each tank to each ecological response, resulting in one GLMM per ecological response. To assess the effects of local genomic diversity, we related the allelic frequencies of each SNP (i.e., 15,799 SNPs) to each ecological response, resulting in 142,191 models. P-values were obtained through permutation tests (1000 permutations) and adjusted for multiple comparisons using the false discovery rate (FDR) correction (p.adjust function, stats package), assuming a threshold of FDR=0.05. The SNPs that were found significantly associated with ecological responses (FDR<0.05) were hereafter termed “candidate SNPs”. The number of candidate SNPs and their identity was recorded for each ecological response.

From these models, we estimated the effect sizes (ES) of both genome-wide and local genomic diversity respectively using the following formula (Nakagawa et al., 2017):

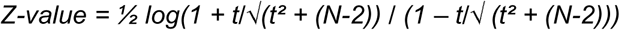

Where *t* is the t-value associated with the parameter of interest (slope of the relationship between genomic diversity and each ecological response) and *N* is the number of observations.

We derived a single Z-value associated with genome-wide diversity (hereafter “genome-wide ES”) for each ecological response. In contrast, local genomic diversity yielded 15,799 Z-values per ecological response, corresponding to one value per SNP. The mean effect size of local genomic diversity (hereafter “local genomic ES”) for each ecological response was calculated as the average absolute Z-value across candidate SNPs only (FDR < 0.05). We then compared genome-wide and local genomic effect sizes across the nine ecological responses using paired *t*-tests, based on absolute Z-values to account for both positive and negative relationships (although most relationships were positive). Because restricting the analysis to candidate SNPs is inherently expected to upwardly bias estimates of local genomic ES, we conducted an additional comparison. Specifically, we performed paired *t*-tests between genome-wide ES calculated only for ecological responses showing significant associations (i.e., total fish biomass, fish biomass variability, and Lymnaeidae abundance; see the Results) and the corresponding local genomic ES for these same responses.

### Is the combined information from a subset of candidate SNPs more impactful than the information of a single candidate SNP?

As we identified that local ES were systematically larger than genome-wide ES (see the Results), we further explored their influence on ecological responses. Specifically, we estimated the ability of the most impactful SNPs to explain ecological responses when considered together (rather than one at a time) to estimate genetic diversity. We expected that combining information from a subset of candidate SNPs into a single metric of genetic diversity (hereafter “candidate SNP subset diversity”) would be more likely to explain ecological responses than the allelic frequency of any single top candidate SNP. If verified, this would suggest an oligogenic architecture for gene-to-ecosystem relationships (rather than monogenic).

To do so, we ranked the SNPs according to their respective ES (from the highest to the lowest) and we estimated the proportion of polymorphic SNPs from the *X* first SNPs, with *X* varying from 2 to 15799 with an increment of 1. This resulted in 15798 incremental estimates of genetic diversity, based on SNPs subsets of varying size (hereafter, “ES_subsetSNPs_”) that we used in 15798 GLMMs similar to that described above. For each ecological response, we identified the size of the SNPs subset that optimize the ES; i.e., *the subset size of SNPs leading to the highest ES for each ecological response*. The size of the optimal subset varied from 3 to 101 SNPs depending on the ecological response (see the Results and Figure 3). We then compared (across ecological responses) these estimates to the ES of the top candidate SNP identified for each ecological response (i.e. the SNP displaying the higher absolute ES, herefter ES_topSNP_) using a paired t-test (based on absolute values). If this resulted into ESsubsetSNPs significantly higher than EStopSNP, this means that combining information from a few markers to estimate genetic diversity of fish populations is more efficient than considering each top candidate SNP individually, and that the ecological impacts of fish is driven by a few SNP rather than a single SNP.

**Figure 3.**
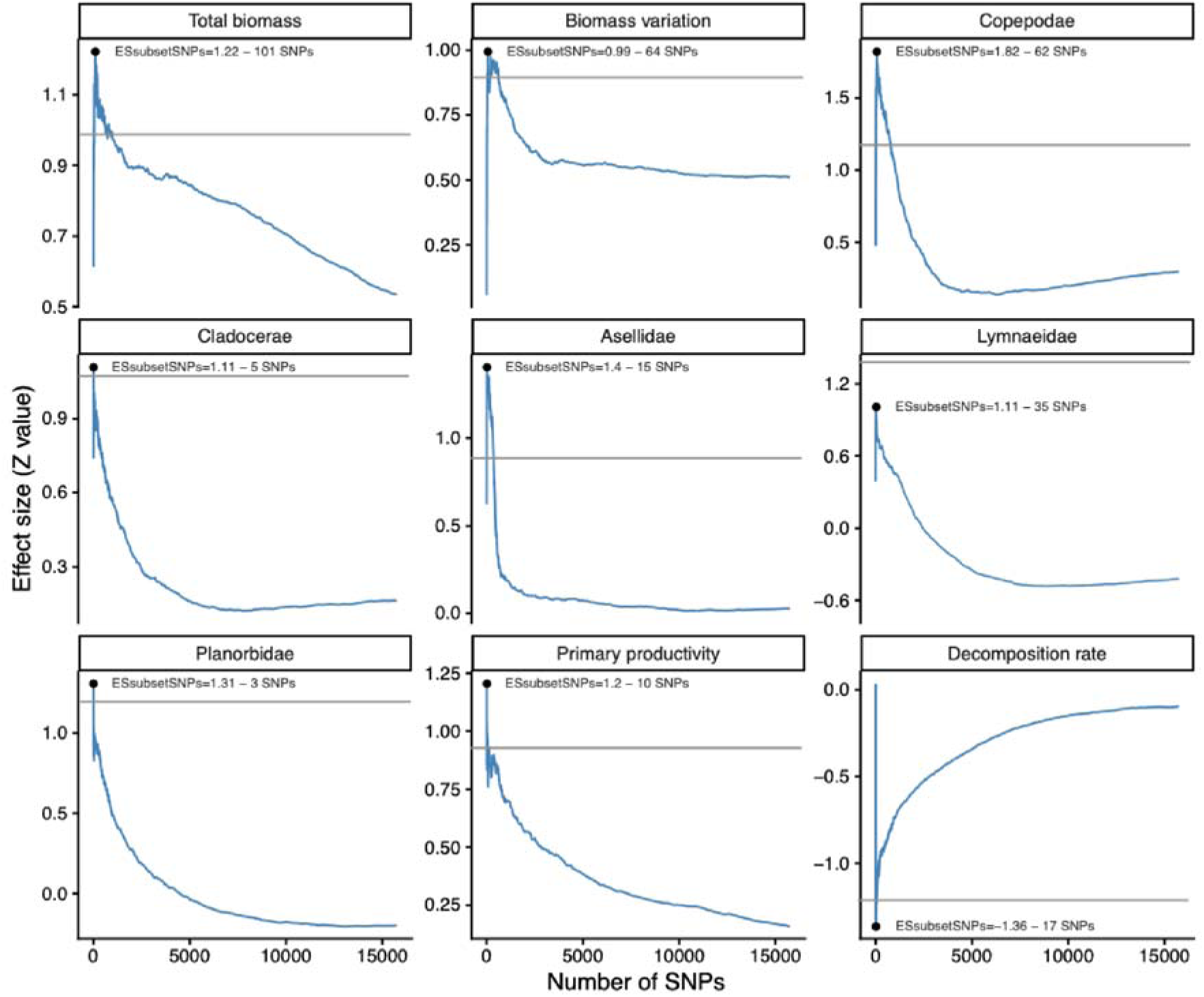
Relationship between the number of candidate SNPs used to estimate genomic diversity of *P. dragarum* (minnow) populations and the effect sizes (Z values) for nine ecological responses. These graphs show the distribution of effect sizes (Z values) measuring the strength of the association between genomic diversity (estimated as the proportion of polymorphic loci) of *P. dragarum* (minnow) populations and each ecological response. Effect sizes were estimated with an iterative process increasing progressively the number of candidate SNPs (initially ranked in decreasing order according to their respective value) used to estimate genomic diversity (from 2 to 15799 candidate SNPs). The distribution of effect sizes is plotted against the number of candidate SNPs used to estimate genomic diversity. For each trait, the black dots indicate the number of candidate SNPs optimizing the effect size (ES_subsetSNPs_). The grey horizontal lines represent the effect size of the top candidate SNP for each ecological response (“ES_topSNP_”, see Figure 1 for exact values). ES_subsetSNPs_ is higher than ES_topSNP_ for all but one ecological response (Lymnaeidae abundance).

To confirm our conclusions, we finally used a model comparison approach (based on the Akaike Information Criteria, AIC) to test whether one of the three metrics (genome-wide diversity, top candidate SNP diversity or candidate SNP subset diversity) is more likely to explain each of the ecological response.

### Comparing global and local genomic ES across climatic conditions

To test the hypothesis that genomic effects on ecological responses are stronger under stressful conditions, we compared local and genome-wide effect sizes (ES) across climatic treatments. We considered the +3°C treatment as a thermal stress for minnows, given their cold-adapted physiology. To this end, we used the same generalized linear mixed models (GLMMs) as described above, but fitted them separately for each climatic condition (i.e., ambient and warmed treatments). For each condition, we estimated both local and genome-wide ES. We then used paired *t*-tests based on absolute Z-values to assess, across ecological responses, whether effect sizes differed between climatic conditions.

### Investigating the functional role of candidate SNPs

To infer the functional role of candidate SNPs associated with ecological responses, we focused on the set of candidate SNPs identified across all ecological responses (n=271, see the Results), assuming that all these SNPs participate to modulating the functioning of the ecosystem (from fish biomass to primary productivity and litter decomposition).

Linkage disequilibrium was first estimated for both the entire set of SNPs and the candidate SNPs to test the hypothesis that LD was not significantly higher in the set of candidate SNPs and that observed results did not arise from clusters of aggregated SNPs. LD was estimated for both sets of SNPs using the SNPRelate R package, LD was calculated as Pearson’s correlation coefficient (r) between pairs of SNPs within a sliding window across the genome. For each SNP pair, LD was expressed as r². The distributions of r² values were summarized using mean values and 95% confidence intervals, allowing comparison of LD patterns between candidate SNPs and the genome-wide background of SNPs.

For functional annotation, we mapped candidate SNP sequences to the *Phoxinus phoxinus* reference genome (GCA_037504875.1^32^) using BLASTN because the *P. dragarum* genome (that we used to genotype the SNPs) was neither chromosome-level assembled nor annotated (note that the *P. phoxinus* genome was not fully annotated). Only high-confidence alignments were retained (percentage identity ≥ 95%, E-value < 1×10⁻⁵), and for each SNP the best-scoring hit was selected based on bit score. SNP genomic coordinates were inferred from the alignment positions, allowing the assignment of SNPs to chromosomes and genomic features. 244 out of the 271 candidate SNPs were successfully mapped to the *P. phoxinus* genome, confirming the phylogenetic similarity of the two species.

Then, coding sequences (CDS) were extracted from the *P. phoxinus* genome annotation and we searched for candidate SNPs overlapping with CDS (findOverlaps functions, GenomicRanges R package). SNPs overlapping annotated CDS were associated with their corresponding gene identifiers, protein products (when available), and strand orientation. To account for potential regulatory variants, SNPs located in promoter regions were also retrieved. Promoters were defined as the 2 kb region upstream of annotated genes and we examined whether some of the candidate SNPs overlapped with such regions. Promoter sequences were extracted from the reference genome, and the nearest coding sequence to each promoter was identified. 49 out of the 244 candidate SNPs were associated either with a CDS (37) or a promoter region (12). We tested whether SNPs associated with ecosystem-level functions were overrepresented in coding sequences or promoters. To do so, we compared the observed number of candidate SNPs overlapping coding regions or promoters (49 out of 244) to a null distribution obtained by randomly sampling the same number of SNPs from the full set of tested SNPs (10,000 permutations). Statistical significance was assessed as the proportion of permutations yielding an equal or greater number of coding SNPs.

Then, CDS harboring candidate SNPs (as well as those close to promoter regions harboring candidate SNPs) were translated into protein sequences and annotated. To do so, proteins corresponding to CDS were extracted from the predicted proteome and aligned against the Swiss-Prot database using BLASTP. Alignments were filtered using stringent criteria (percentage identity ≥ 70%, E-value < 1×10⁻¹⁰), and for each query protein the best hit was retained by prioritizing *Danio rerio* annotations when available (the closest annotated genome). When not available, orthologous proteins from well-annotated vertebrates (human, mouse…) were retained (based on bit score) to maximize functional inference. UniProt accession numbers were extracted from Swiss-Prot identifiers and used as functional proxies for downstream analyses. Gene Ontology (GO) terms and pathway annotations were parsed from Swiss-Prot annotation fields when available.

A GO enrichment analysis was then performed to identify the most likely biological functions involved in the ecological impacts of *P. dragarum*. Because the number of SNPs associated with a CDS or a promoter for which we identified a protein name was small (n=21), ontology enrichment was performed by combining all these SNPs to increase statistical power. Enrichment of Biological Process (BP) GO terms was assessed using the enrichGO R function (clusterProfiler R package), with the *Danio rerio* annotation database (org.Dr.eg.db) used as a reference due to the absence of a species-specific annotation resource. UniProt identifiers were used as gene identifiers, and enrichment significance was evaluated using Benjamini–Hochberg–corrected p-values, with a q-value threshold of 0.05. Enriched GO terms were summarized based on fold enrichment, gene counts, and adjusted p-values.

## RESULTS

### Local genomic effects are stronger than genome-wide effects

We found that population genetic diversity measured at specific candidate SNPs was more strongly associated with ecological responses than genome-wide diversity (paired t-test, t=-8.167, d.f.=8, p<0.001, Figure 1), and that these candidate SNPs had strong and consistent effects on ecological responses spanning from population to ecosystem responses. Indeed, significant candidate SNPs were identified for all ecological responses (from 2 to 95 significant candidate SNPs per variable, Figure 1A), and they were all very strongly associated with their respective ecological response (mean local genomic ES=0.906, 95%CI=0.852-0.959). For instance, the top candidate SNP identified for each trait (i.e., the SNP having the strongest ES for a given trait, “ES_topSNP_”) reached an effect size ranging from 0.90 to 1.34 depending on the trait (corresponding to 0.717<r_Pearson_<0.871, see Figure 1 and Figure 2d-f for some illustration). Contrastingly, genome-wide ES were significant for only 3 out of the 9 ecological responses (total fish biomass, fish biomass variation and abundance of Lymanaeidae, Figure 1A), and they were weak to moderate in strength (mean genome-wide ES=0.262, 95%CI=0.138-0.386, Figure 1 and Figure 2a-c for some illustration, Table S1 for values). Noteworthily, genome-wide ES were poorly influenced by the metric used to estimate genome-wide diversity since we obtained similar results using either the expected heterozygosity or a genetic distance based on the number of SNPs by which two individuals differ (Table S1).

The same conclusion was also reached when only the three significant genome-wide ES were selected and compared to their local counterparts (paired t-test, t=-6.921, d.f.=2, p=0.020). In this case, local genomic ES (mean local genomic ES=0.865, 95%CI=0.788-0.941, n=3) was almost twice higher in absolute value than genome-wide ES (mean genome-wide ES=0.493. 95%CI=0.463-0.521, n=3).

### Genomic diversity estimated from a subset of candidate SNPs is more impactful than diversity estimated from a single candidate SNP

Genomic ES estimated from a subset of candidate SNPs (the size of the subset varying from 3 to 101 SNPs depending on the ecological response, see Figure 3) tended to be stronger than local genomic ES estimated from the top candidate SNP of each trait (paired t-test, t=-2.160, d.f.=8, p=0.068, see Figure 2g-i for some illustration). Indeed, for all ecological responses (but Lymnaeidae abundance), ES_subsetSNPs_ was stronger than Es_topSNP_ (Figure 3).

### The impacts of fish on ecological responses may be oligogenic

Using a model comparison approach, we confirm that candidate SNP subset diversity outperformed both genome-wide diversity and top candidate SNP diversity in explaining each ecological responses (ΔAIC>4 for all comparisons, see Table S2). Hence, genetic diversity estimated from a restricted set of candidate SNPs provided a better explanation of ecological variation than either single-locus variation or genome-wide diversity. This pattern suggests that gene-to-ecosystem relationships may follow an oligogenic architecture, where a limited number of SNPs jointly contribute to ecosystem-level variation, rather than a purely monogenic or a diffuse polygenic architecture.

### Global and local genomic ES are similar across climatic conditions

Models performed for the two climatic treatments separately revealed that global genomic ES were not significantly different between the warm (more stressful) and the ambient climate (paired t-test for absolute ES values, t=-1.489, d.f.=8, p=0.175; p=0.126 for raw ES values). Similarly, local genomic ES were not significantly different between the warm and the ambient climate, when all variable responses were considered (paired t-test for absolute ES values, t=-1.319, d.f.=8, p=0.223; p=0.541 for raw ES values). This suggests that effect sizes (measured either at the local or global genomic scale) were relatively similar and consistent across two contrasting environmental conditions.

### The impacts of fish on ecological responses are sustained by a few functional pathways

A total of 271 SNPs were identified as candidates for at least one of the nine ecological response variables, including nine SNPs associated with two responses. Candidate SNPs were distributed across the 25 chromosomes of *P. dragarum*, with the number of SNPs per chromosome ranging from 4 (chromosome 12) to 21 (chromosome 1), and there was no evidence for a strong clustering in a particular genomic region (Figure S1). The number of candidate SNPs per chromosome was not significantly associated with the chromosome size, although a positive trend was observed (Figure S2). The level of linkage disequilibrium was low, and candidate SNPs exhibited lower levels of linkage disequilibrium compared to the genome-wide background; the mean LD (r²) among candidate SNPs was 0.0052 (95% CI: 0.0051–0.0053), whereas the mean LD across all SNPs was 0.0088 (95% CI: 0.00875–0.00880). The non-overlapping confidence intervals indicate that candidate SNPs are significantly less correlated than expected from genome-wide LD patterns. Among these 271 SNPs, 37 were located within coding DNA sequences (CDS) and 12 within promoter regions of CDS (Table S3). This proportion was significantly higher than expected by chance, as assessed by a permutation test (expected mean = 29.22 ± 4.98 SD; p < 0.001).

Protein annotations could be assigned to 21 of the 49 SNPs located within CDS or promoter regions (Table S3). The corresponding annotated genes were used for Gene Ontology (GO) enrichment analysis, which identified 56 significantly enriched Biological Process (BP) terms (FDR < 0.05; Table S4). Although individual GO terms spanned diverse biological processes, they converged into five broad functional axes underlying the ecological effects of *P. dragarum*. These axes encompassed: (1) multicellular development and organization, supporting organism growth and biomass accumulation; (2) cellular differentiation and tissue specialization, contributing to functional plasticity; (3) physiological support systems, including cardiovascular and metabolic processes essential for organism performance; (4) protein metabolism and post-translational regulation, shaping energy allocation, metabolic efficiency and adaptive responses; and (5) regulation of growth and biological processes, integrating environmental signals to modulate growth and ecosystem-level responses (Table 1). Strikingly, these functional axes were driven by a small set of ecologically important genes (amotl2a, clvs2, dip2ba, krt8, and ccdc69; Table 1, Table S4). Through likely pleiotropic effects, these genes collectively mediated the influence of *P. dragarum* on fish biomass, prey biomass and multiple ecosystem functions.

## DISCUSSION

Using an ecologically-realistic mesocosm experiment spanning multiple trophic levels, we demonstrate that genetic variation at a few important gene regions in a top-predatory fish can alter ecosystem processes, from prey abundances to ecosystem functioning. This finding reinforces growing evidence that intraspecific genetic variation can act as a substantial driver of ecosystem dynamics^4,5,13,14^, and provides one of the most convincing demonstrations to date that gene-to-ecosystem relationships are sustained by a restricted part of the genome, rather than by genome-wide variation. This pattern aligns with the hypothesis that a small subset of genes contributes to ecosystem-level variation^18^. However, our results are consistent with a small set of gene regions representing an oligogenic architecture of traits underlying ecological effects, rather than a monogenic architecture, as implicitly suggested by the “keystone genes” framework^18^. Notably, SNPs linked to these gene regions exhibited strong and consistent effects across contrasting climatic conditions, suggesting a robust signal linking intraspecific genetic variation to ecosystem processes. Importantly, we further identify candidate functional pathways underlying these relationships, thus providing a putative mechanistic basis for linking intraspecific diversity, energy use, and trophic transfer across trophic levels.

A central result of our study is that the ecological consequences of intraspecific genetic variation are not evenly distributed across the genome. Within populations, ecological complementarity among individuals is expected to enhance resource use efficiency and energy transfer across trophic levels^14,33^. Complementarity is often assumed to arise from phenotypic, behavioural, and physiological differences among individuals partly structured by genetic variation^13,34^. Genome-wide diversity has therefore frequently been used as a proxy for such functional differences among individuals. However, we found that genome-wide diversity explained relatively few ecological responses with weak to moderate effect sizes, yet comparable to those reported in a previous meta-analysis^5^. This moderate coupling between genome-wide diversity and ecological responses challenges the assumption that genomic effects scale with genome-wide diversity. This might partly be explained by the fact that most parts of the genome are non-functional (i.e., do not code for phenotypic traits) and may actually blur the effects of functional genomic diversity. Accordingly, ecological responses were strongly associated with a restricted set of SNPs (i.e., candidate SNPs) that were -for some of them- associated with essential biological functions^13,18,19^. Although a few studies have identified strong links between specific genes and ecological processes, these have often been conducted in simplified or highly controlled systems, frequently in plant models such as *Aribidopsis*^19,21,23,24^. Here, we extend this evidence to a more complex and realistic multitrophic context, showing that such genomic signals can propagate across multiple trophic levels, from fish biomass to prey communities and to ecosystem processes such as primary productivity and organic matter decomposition.

We further found that these genomic effects were consistent across ambient and warm conditions (+3°C), suggesting that, in this study, the genomic architecture of traits underlying ecological effects was relatively robust to environmental variation within the tested range. The warming scenario that we set follows realistic and predicted climatic scenarios (a +3 °C treatment reflects the magnitude of warming projected by the end of the 21st century under SSP2-4.5 to SSP5-8.5 scenarios^35^), which suggests that in some cases climate change may influence ecosystems independently from changes in the genomic diversity of populations^28^. This environmental stability contrasts with expectations from strong genotype-by-environment interactions often reported in eco-evolutionary systems^36^, and suggests that genomic drivers of ecological responses may persist despite realistic environmental changes, demonstrating their importance for ecosystem functioning.

Although we identified candidate SNPs with strong effect sizes across all ecological responses, we found that quantifying genetic diversity from a subset of 3 to 101 of these candidate SNPs systematically outperformed both single-locus and genome-wide estimates of genetic diversity. Although this finding requires additional works to establish and confirm the mechanistic associations between candidate gene regions, phenotypic traits and their influence on resources, this result is consistent with an oligogenic architecture of gene-to-ecosystem relationships, where a limited number of loci jointly influence traits that scale up to ecological processes. Such architectures occupy an intermediate position between classical large-effect “keystone gene” models^18^ (i.e., a monogenic architecture) and highly polygenic (omnigenic) frameworks^37^, and align with empirical evidence that moderate numbers of loci can have disproportionate effects on complex traits^37,38^. Up-to-our knowledge, most previous studies having established significant links between genes and ecosystem processes have invoked or suggested monogenic architectures^18,19,21–24^, which, overall, suggests that the genomic architecture of gene-to-ecosystem relationships might actually span a continuum from single-locus to multi-locus architecture. Yet, a striking result of our study is that the small subsets of loci we identified have extremely strong explanatory powers for all ecological responses, suggesting that these subsets of gene regions associated with candidate loci are relevant and strong predictors for ecological dynamics in realistic ecosystems.

We further provide evidence for the functional bases underlying these gene-to-ecosystem relationships by revealing that candidate SNPs are enriched in biological processes related to growth, development, and metabolic regulation. These pathways are central to energy allocation and organism performance, which are directly associated with food acquisition and energy fluxes among trophic levels. Although more precise genetic-phenotypic linkage are needed to reveal more precisely these mechanistic pathways, this finding is coherent since such pathways indirectly mediate trophic interactions and hence ecosystem processes^39,40^. Moreover, although our study is based on a reduced-genome representation and a non-model organism (for which relatively few functional studies have been conducted), we identified a few specific gene regions associated with the biological processes cited above (amotl2a, clvs2, dip2ba, krt8, and ccdc69). Several of these gene regions have pleiotropic effects and are conserved across phyla. For instance, the krt8 gene codes for a major protein in the cytoskeleton of fish, that has been shown to be associated to functions ranging from cell migration and proliferation (cell development) to immune regulation^41^, and is largely conserved across the Vertebra phyla^42^.

At this stage, we can only speculate regarding the linkages between these gene regions, associated biological processes, and the phenotypic traits sustaining the ecosystem impacts of minnows. Hereafter, we provide a hypothetical causal explanation for the gene-to-ecosystem relationships identified in this study. We previously revealed striking phenotypic differences among some of the populations used to seed the mesocosms; the general activity, the adult body mass and the resting metabolic rate varied substantially among minnow populations^27^. In fish, these traits are strongly associated with the consumption of prey resources; for instance, larger individuals can consume larger prey (due to gap-size limitation in smaller fish), whereas more active fish generally consume more prey per unit of time. Accordingly, we found that these traits discriminating minnow populations were significantly associated with ecosystem responses; for instance, minnow adult body size was associated with the abundance of prey, whereas resting metabolic rate was associated with primary productivity^27^. It is therefore possible that some candidate gene regions identified in this study are controlling (at least partly) some of the traits discriminating minnow populations and altering ecosystems responses; increased polymorphism at these gene regions (i.e., diversity at the subset of candidate SNPs) increases phenotypic variance in minnow populations (and hence complementarity) and optimizes resource use by limiting niche overlap among individuals. Future studies should improve our understanding of the links between genes, phenotypic traits and ecosystem dynamics (beyond model species such as *Arabidopsis*) by combining controlled experiments, comparative and functional genomics, and long-term ecological monitoring.

To conclude, our results indicate that gene-to-ecosystem relationships depend more strongly on the functional genomic distribution of trait variation than on its genome-wide magnitude. This suggests a need to move beyond genome-wide summaries of genetic diversity toward functionally informed genomic perspectives that explicitly account for the architecture of ecologically-important traits^43^. Such approach may permit identifying pleiotropic and conserved genes substantially associated with ecosystem dynamics (i.e., sustaining gene-to-ecosystem relationships) in a large set of non-model species, which might be particularly relevant for estimating functional genomic diversity at the *community level* (i.e., beyond a focal-species approach), and better grasp the reciprocal dynamics between biodiversity (measured along the intra- and interspecific continuum) changes and ecosystem functioning^see, 20^. Moreover, our findings suggest that ecological dynamics in complex ecosystems could be predicted based on estimates of genetic diversity measured from a very few functional genes; this opens fascinating perspectives for predicting the ecological consequences of intraspecific biodiversity loss. By identifying a restricted set of SNPs associated with ecosystem-level processes, our study hence provides a step toward predictive eco-evolutionary frameworks. Rather than relying on genome-wide diversity metrics, future models could incorporate functionally relevant genomic variation to predict ecosystem responses to environmental change^20^. However, translating such approaches to natural populations remains challenging, particularly due to environmental heterogeneity and the difficulty of identifying ecologically relevant genes across systems.

## ACKNOWLEDGMENTS

We warmly thank all the colleagues who contributed to the experiment and who provided comments on this work.

## DATA AVAILABILITY

The primary and secondary data used in this MS, as well as R scripts, are available at https://doi.org/10.6084/m9.figshare.33965854

## AUTHOR CONTRIBUTION

**Maxime Stanislawek:** Software (Lead); Validation (Lead); Formal analysis (Lead); Investigation (Lead); Data curation (Lead); Writing - original draft (Lead); Writing - review & editing (Equal); Visualization (Lead); **Patrik Nosil:** Methodology (Supporting); Software (Supporting); Validation (Lead); Formal analysis (Supporting); Writing - original draft (Supporting); Writing - review & editing (Equal); **Allan Raffard:** Conceptualization (Equal); Methodology (Equal); Software (Lead); Validation (Lead); Formal analysis (Supporting); Investigation (Equal); Writing - original draft (Lead); Writing - review & editing (Supporting); Visualization (Supporting); **Laura Fargeot:** Methodology (Equal); Data curation (Supporting); Writing - original draft (Supporting); Writing - review & editing (Supporting); Visualization (Supporting); **Murielle Richard:** Conceptualization (Supporting); Methodology (Supporting); Validation (Supporting); Investigation (Supporting); Data curation (Supporting); Writing - original draft (Supporting); Writing - review & editing (Supporting); Visualization (Supporting); Supervision (Supporting; **Madoka Krick:** Methodology (Supporting); Software (Supporting); Formal analysis (Supporting); Resources (Supporting); Data curation (Supporting); Writing - original draft (Supporting); Writing - review & editing (Supporting); **Camille Poesy:** Methodology (Supporting); Software (Supporting); Formal analysis (Supporting); Resources (Supporting); Data curation (Supporting); Writing - original draft (Supporting); Writing - review & editing (Supporting); **Maxim Lefort:** Methodology (Supporting); Software (Supporting); Formal analysis (Supporting); Resources (Supporting); Data curation (Supporting); Writing - original draft (Supporting); Writing - review & editing (Supporting); **Geraldine Loot:** Conceptualization (Supporting); Methodology (Supporting; Validation (Supporting); Resources (Supporting); Writing - original draft (Supporting); Writing - review & editing (Supporting); Supervision (Supporting); **Simon Blanchet:** Conceptualization (Lead); Methodology (Lead); Software (Supporting); Validation (Lead); Formal analysis (Supporting); Investigation (Lead); Resources (Equal); Data curation (Supporting); Writing - original draft (Equal); Writing - review & editing (Lead); Visualization (Lead); Supervision (Lead); Project administration (Lead); Funding acquisition (Lead).

## Supplementary material

**Figure S1.**
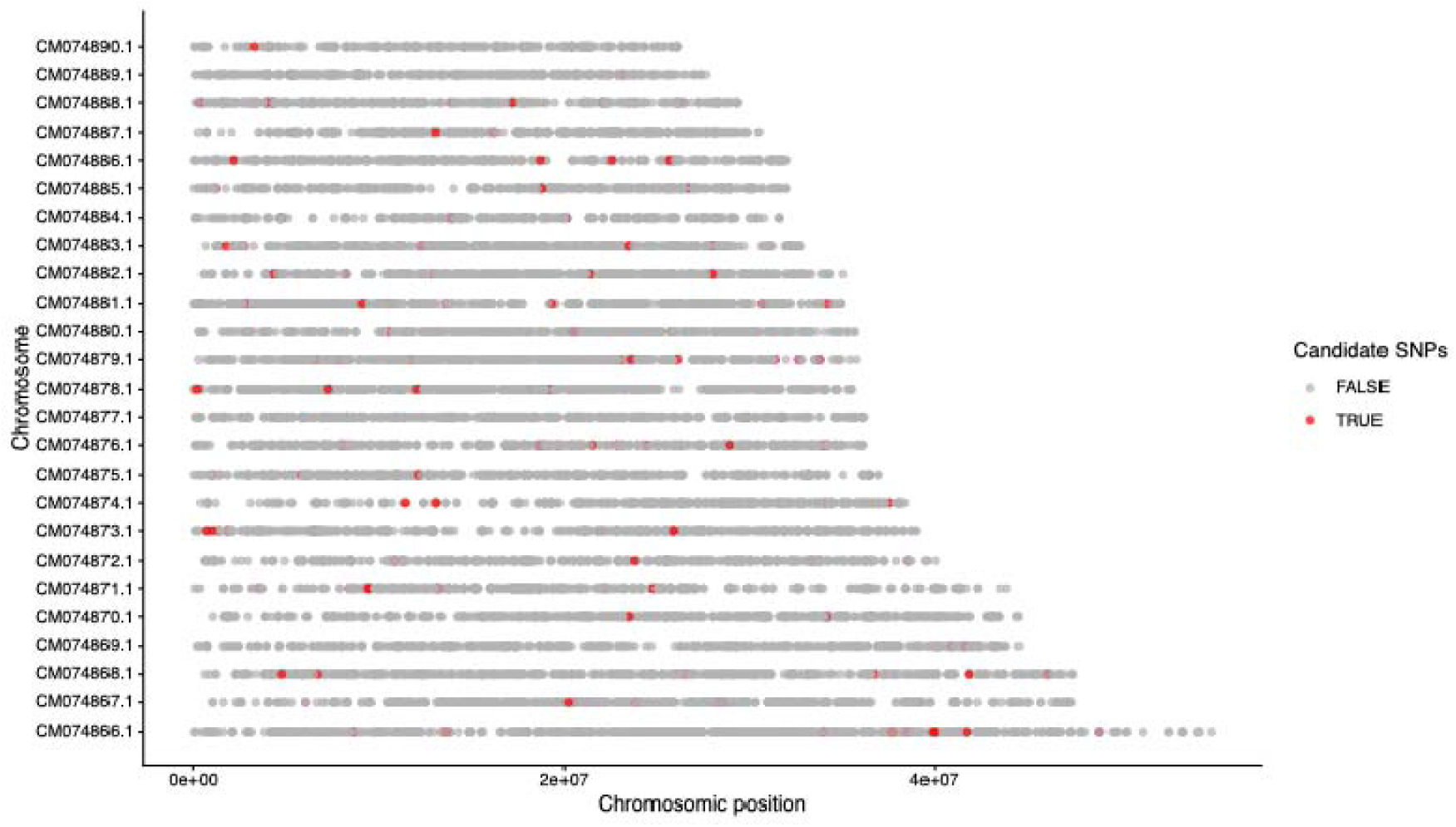
Distribution of the candidate SNPs (n=274, red dots) along each of the chromosome of *Phoxinus phoxinus*. The mapping and position of each SNP to the Phoxinus genome has been done using the *P. phoxinus* reference genome rather than the *P. dragarum* genome as the former was better annotated.

**Figure S2.**
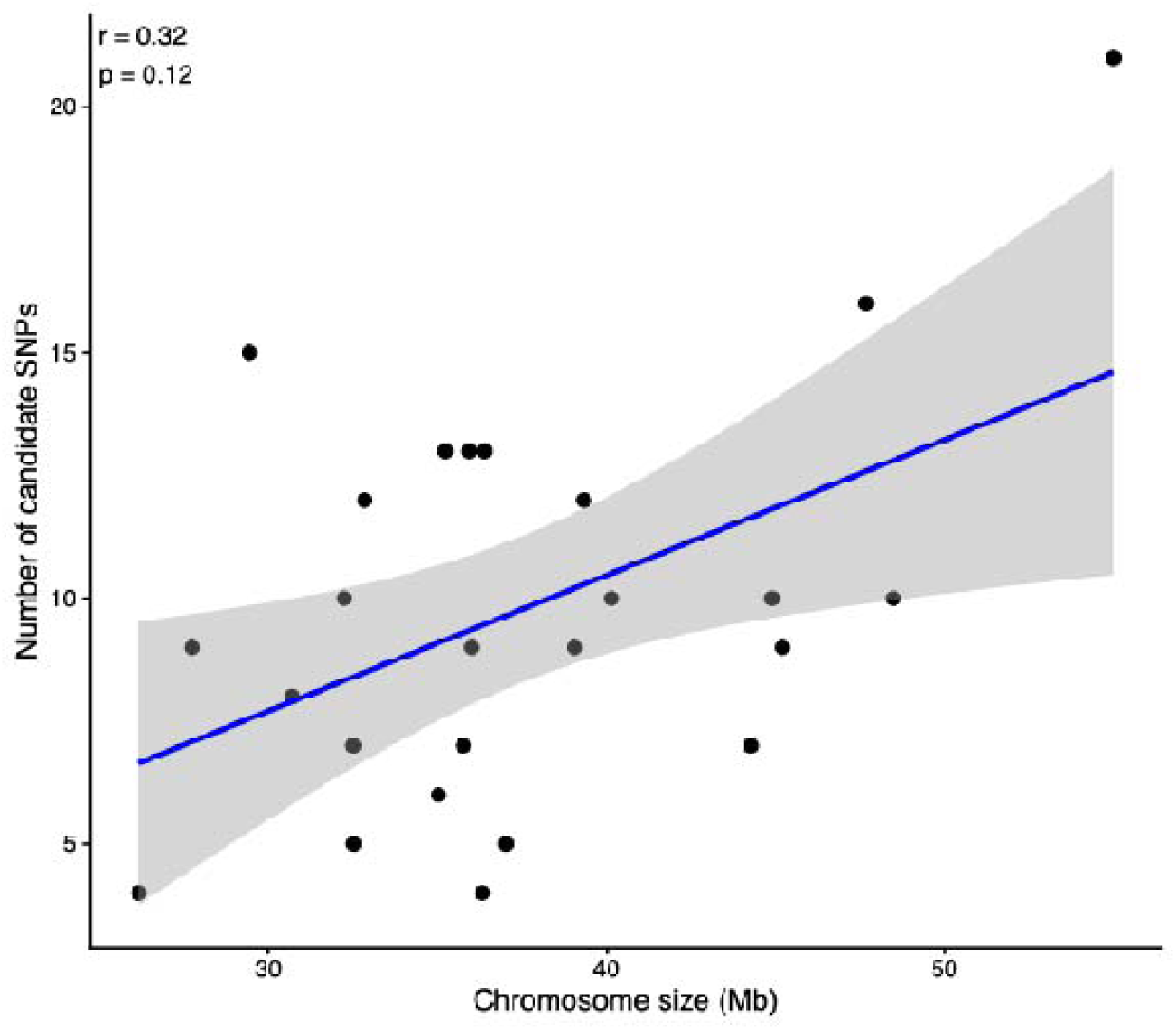
Relationship between the number of candidate SNP per chromosome and the size (expressed as the number of bases) of each chromosome. Chromosome size was estimated from the annotate *Phoxinus phoxinus* genome. The Pearson coefficient of correlation is provided, as well as its associated p-value. Each dot represents a chromosome.

**Table S1.**
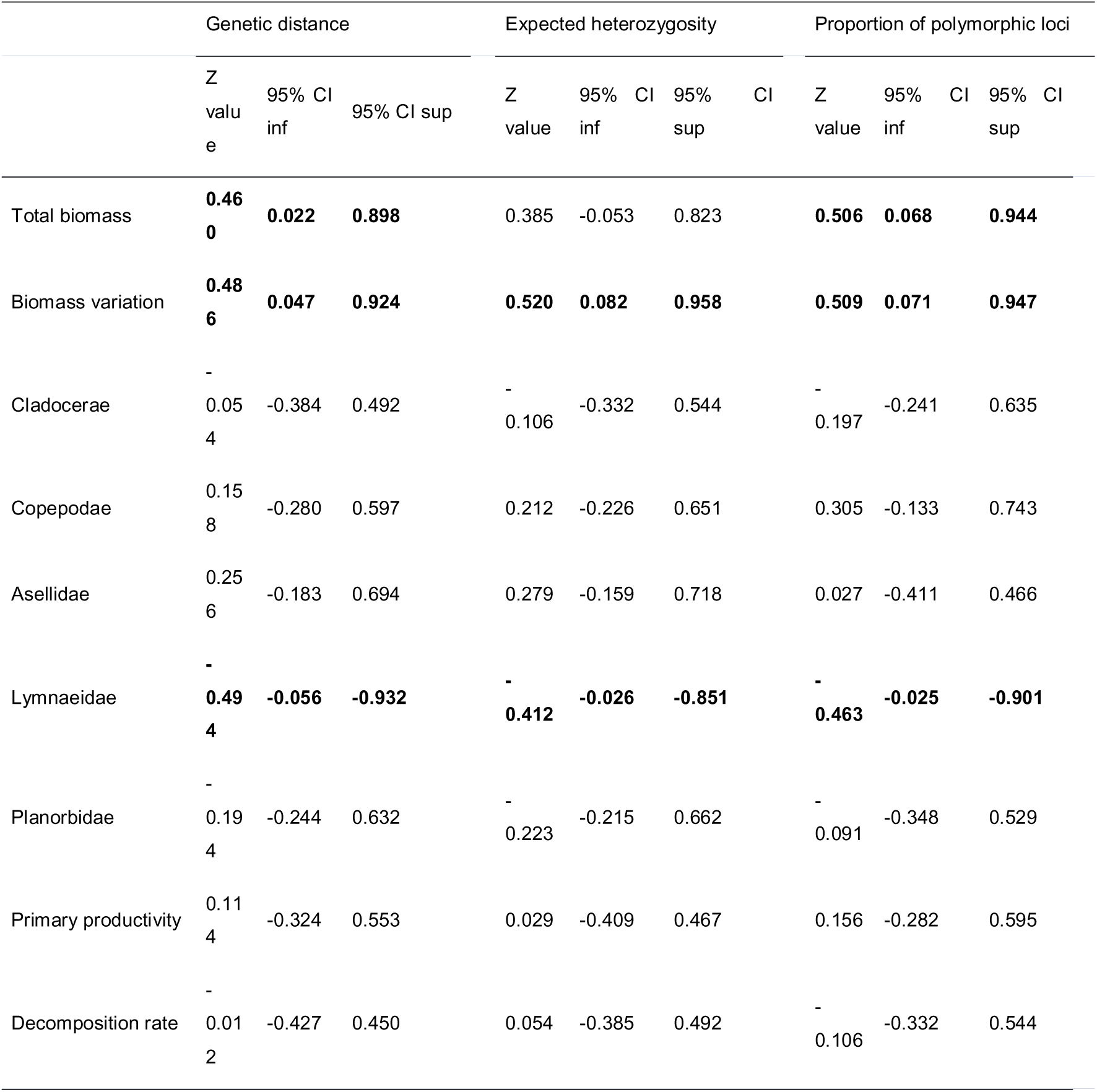
Comparison of genome-wide genomic diversity metrics for predicting ecological responses. Genome-wide genomic effects were estimated using three alternative measures of genomic diversity: a genetic distance based on the number of loci by which two individuals differ, the expected heterozygosity, and the proportion of polymorphic SNPs. For each ecological response, the table reports the standardized effect size (Z-value) and associated 95% confidence intervals obtained from generalized linear mixed models. Similar effect sizes across diversity metrics indicate that genome-wide genomic effects are robust to the choice of diversity estimator. Bold values indicate significant estimates (95%CIs do not overlap 0).

**Table S2.** Comparison of alternative genomic diversity metrics for explaining ecological responses. Generalized linear mixed models relating ecological responses to three measures of genomic diversity were compared using Akaike Information Criteria (AIC). The table reports the AIC values for models based on genomic diversity estimated from the optimal subset of candidate SNPs (“Candidate SNP subset diversity”), the most strongly associated single candidate SNP (“Top candidate SNP diversity”), and genome-wide diversity. Lower AIC values indicate stronger support for the corresponding model. For all ecological responses, the “best SNP subset diversity” metric had the stronger support, with differences in AIC values (ΔAIC) always falling above 4 units for all responses but the biomass variation, which indicates strong support for the best model.

|  | Total<br>biomass | Biomass<br>variation | Copepodae | Cladocerae | Asellidae | Lymnaeidae | Planorbidae | Primary<br>productivity | Decomposition<br>rate |
| --- | --- | --- | --- | --- | --- | --- | --- | --- | --- |
| Candidate<br>SNP subset<br>diversity | **159.4** | **-56.4** | **243.5** | **116.1** | **177.8** | **243.5** | **79.8** | **224.5** | **-300.7** |
| Top candidate<br>SNP diversity | 165.0 | -53.3 | 253.0 | 120.9 | 183.6 | 253.0 | 87.7 | 249.0 | -290.6 |
| Genome-wide<br>diversity | 180.8 | -42.4 | 258.1 | 121.9 | 186.7 | 259.3 | 97.9 | 251.4 | -269.7 |

**Table S3.**
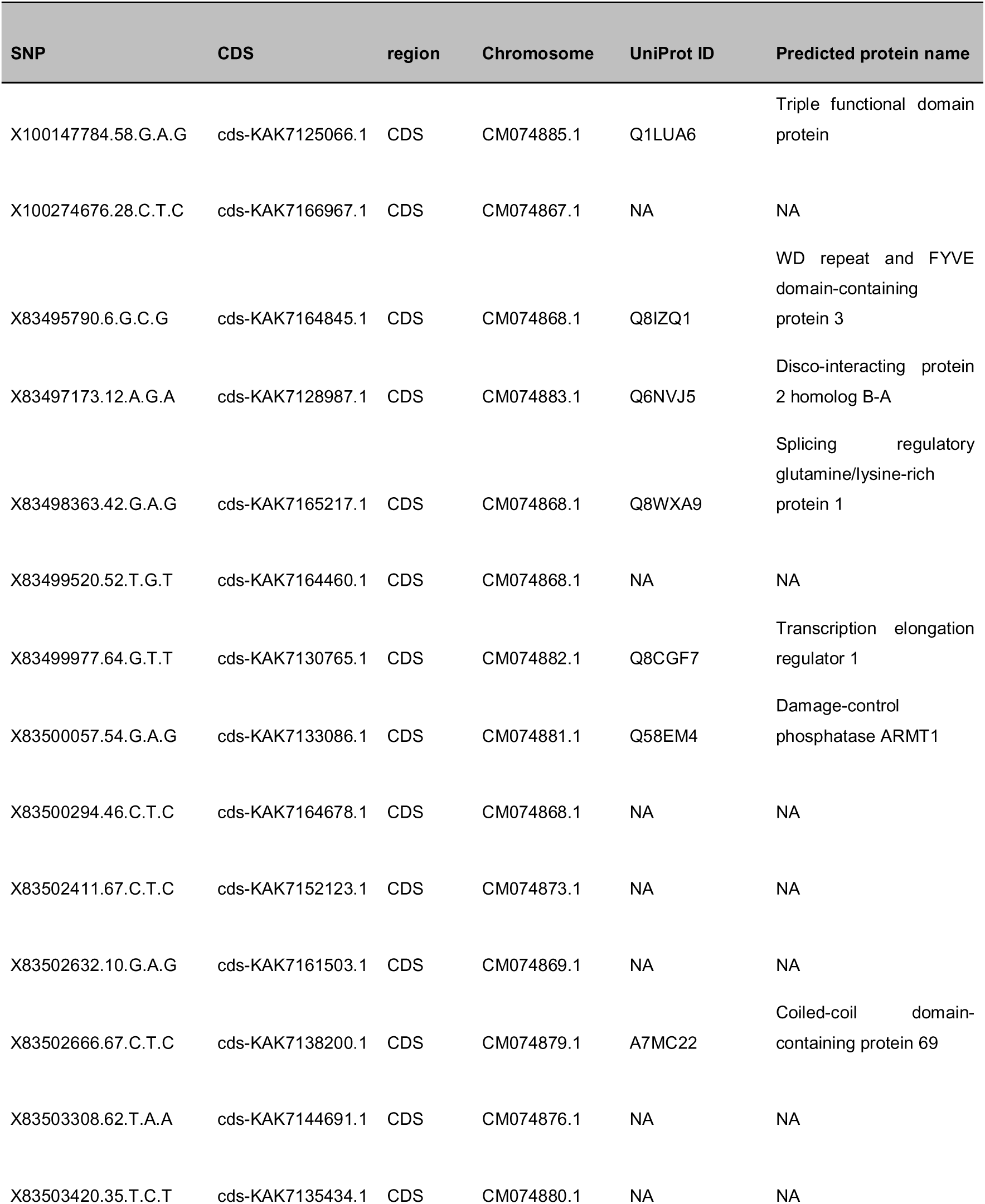
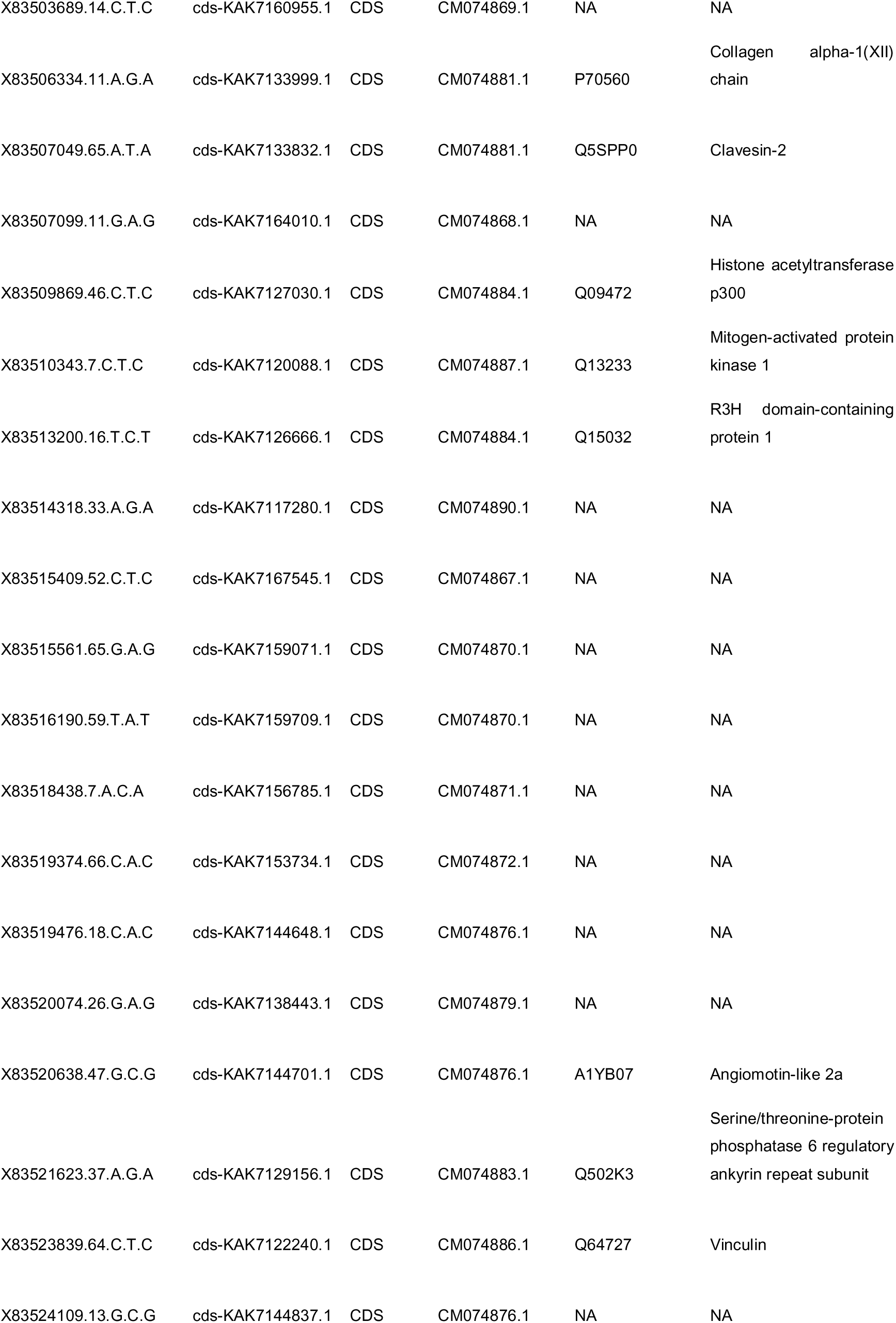
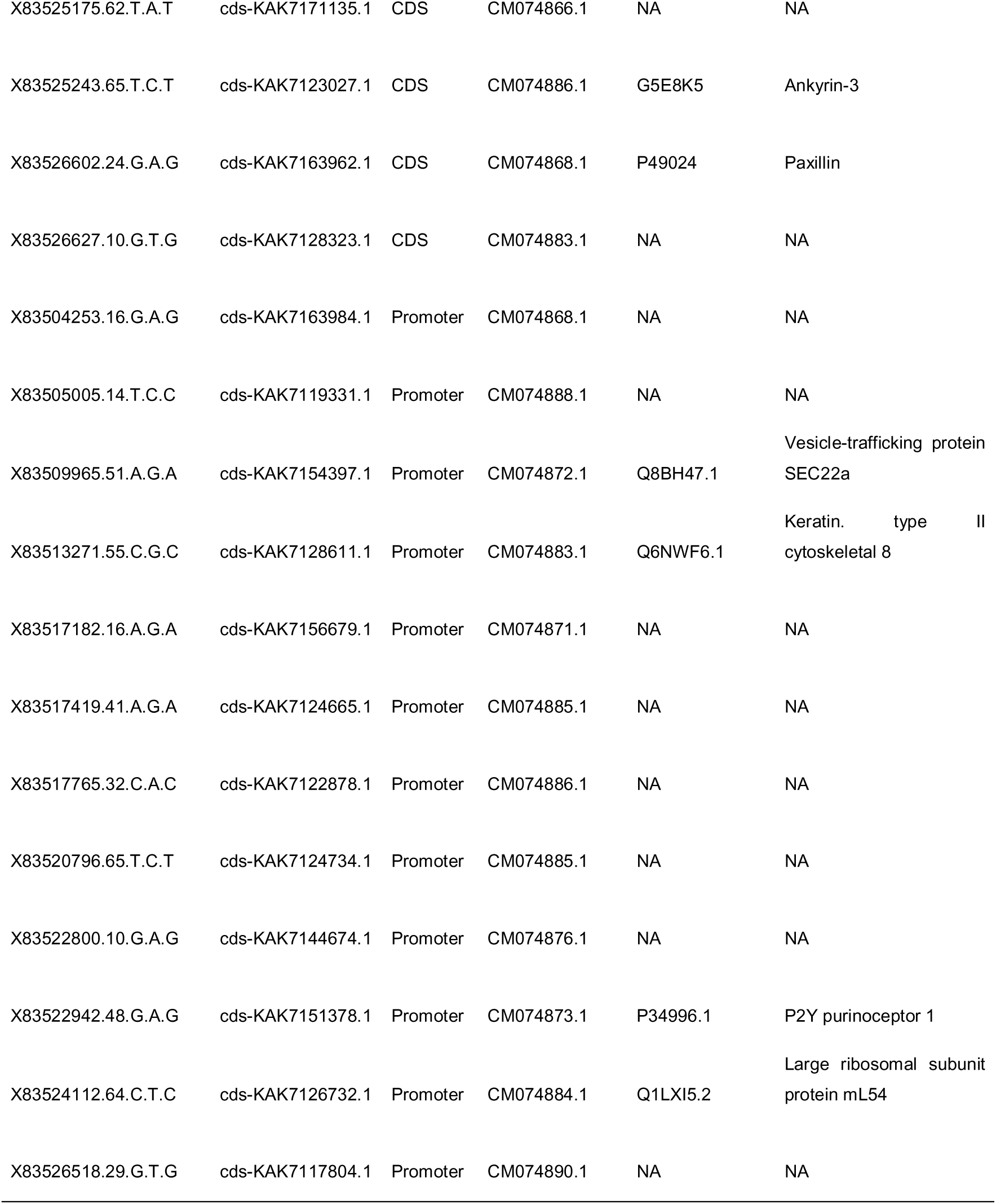
Functional annotation of candidate SNPs associated with ecological responses. The table reports the 49 candidate SNPs located within coding DNA sequences (CDS) or promoter regions after mapping to the annotated *Phoxinus phoxinus* reference genome. For each SNP, the corresponding genomic feature, chromosome, coding sequence (CDS) identifier, UniProt accession number, and predicted protein are provided when a significant protein annotation could be assigned. Entries marked “NA” indicate that no reliable protein annotation could be assigned (n=28 out of 49).

| SNP | CDS | region | Chromosome | UniProt ID | Predicted protein name |
| --- | --- | --- | --- | --- | --- |
| X100147784.58.G.A.G | cds-KAK7125066.1 | CDS | CM074885.1 | Q1LUA6 | Triple functional domain protein |
| X100274676.28.C.T.C | cds-KAK7166967.1 | CDS | CM074867.1 | NA | NA |
| X83495790.6.G.C.G | cds-KAK7164845.1 | CDS | CM074868.1 | Q8IZQ1 | WD repeat and FYVE domain-containing protein 3 |
| X83497173.12.A.G.A | cds-KAK7128987.1 | CDS | CM074883.1 | Q6NVJ5 | Disco-interacting protein 2 homolog B-A |
| X83498363.42.G.A.G | cds-KAK7165217.1 | CDS | CM074868.1 | Q8WXA9 | Splicing regulatory glutamine/lysine-rich protein 1 |
| X83499520.52.T.G.T | cds-KAK7164460.1 | CDS | CM074868.1 | NA | NA |
| X83499977.64.G.T.T | cds-KAK7130765.1 | CDS | CM074882.1 | Q8CGF7 | Transcription elongation regulator 1 |
| X83500057.54.G.A.G | cds-KAK7133086.1 | CDS | CM074881.1 | Q58EM4 | Damage-control phosphatase ARMT1 |
| X83500294.46.C.T.C | cds-KAK7164678.1 | CDS | CM074868.1 | NA | NA |
| X83502411.67.C.T.C | cds-KAK7152123.1 | CDS | CM074873.1 | NA | NA |
| X83502632.10.G.A.G | cds-KAK7161503.1 | CDS | CM074869.1 | NA | NA |
| X83502666.67.C.T.C | cds-KAK7138200.1 | CDS | CM074879.1 | A7MC22 | Coiled-coil domain-containing protein 69 |
| X83503308.62.T.A.A | cds-KAK7144691.1 | CDS | CM074876.1 | NA | NA |
| X83503420.35.T.C.T | cds-KAK7135434.1 | CDS | CM074880.1 | NA | NA |
| X83503689.14.C.T.C | cds-KAK7160955.1 | CDS | CM074869.1 | NA | NA |
| X83506334.11.A.G.A | cds-KAK7133999.1 | CDS | CM074881.1 | P70560 | Collagen alpha-1(XII) chain |
| X83507049.65.A.T.A | cds-KAK7133832.1 | CDS | CM074881.1 | Q5SPP0 | Clavesin-2 |
| X83507099.11.G.A.G | cds-KAK7164010.1 | CDS | CM074868.1 | NA | NA |
| X83509869.46.C.T.C | cds-KAK7127030.1 | CDS | CM074884.1 | Q09472 | Histone acetyltransferase p300 |
| X83510343.7.C.T.C | cds-KAK7120088.1 | CDS | CM074887.1 | Q13233 | Mitogen-activated protein kinase 1 |
| X83513200.16.T.C.T | cds-KAK7126666.1 | CDS | CM074884.1 | Q15032 | R3H domain-containing protein 1 |
| X83514318.33.A.G.A | cds-KAK7117280.1 | CDS | CM074890.1 | NA | NA |
| X83515409.52.C.T.C | cds-KAK7167545.1 | CDS | CM074867.1 | NA | NA |
| X83515561.65.G.A.G | cds-KAK7159071.1 | CDS | CM074870.1 | NA | NA |
| X83516190.59.T.A.T | cds-KAK7159709.1 | CDS | CM074870.1 | NA | NA |
| X83518438.7.A.C.A | cds-KAK7156785.1 | CDS | CM074871.1 | NA | NA |
| X83519374.66.C.A.C | cds-KAK7153734.1 | CDS | CM074872.1 | NA | NA |
| X83519476.18.C.A.C | cds-KAK7144648.1 | CDS | CM074876.1 | NA | NA |
| X83520074.26.G.A.G | cds-KAK7138443.1 | CDS | CM074879.1 | NA | NA |
| X83520638.47.G.C.G | cds-KAK7144701.1 | CDS | CM074876.1 | A1YB07 | Angiomotin-like 2a |
| X83521623.37.A.G.A | cds-KAK7129156.1 | CDS | CM074883.1 | Q502K3 | Serine/threonine-protein phosphatase 6 regulatory ankyrin repeat subunit |
| X83523839.64.C.T.C | cds-KAK7122240.1 | CDS | CM074886.1 | Q64727 | Vinculin |
| X83524109.13.G.C.G | cds-KAK7144837.1 | CDS | CM074876.1 | NA | NA |
| X83525175.62.T.A.T | cds-KAK7171135.1 | CDS | CM074866.1 | NA | NA |
| X83525243.65.T.C.T | cds-KAK7123027.1 | CDS | CM074886.1 | G5E8K5 | Ankyrin-3 |
| X83526602.24.G.A.G | cds-KAK7163962.1 | CDS | CM074868.1 | P49024 | Paxillin |
| X83526627.10.G.T.G | cds-KAK7128323.1 | CDS | CM074883.1 | NA | NA |
| X83504253.16.G.A.G | cds-KAK7163984.1 | Promoter | CM074868.1 | NA | NA |
| X83505005.14.T.C.C | cds-KAK7119331.1 | Promoter | CM074888.1 | NA | NA |
| X83509965.51.A.G.A | cds-KAK7154397.1 | Promoter | CM074872.1 | Q8BH47.1 | Vesicle-trafficking protein<br>SEC22a |
| X83513271.55.C.G.C | cds-KAK7128611.1 | Promoter | CM074883.1 | Q6NWF6.1 | Keratin. type II<br>cytoskeletal 8 |
| X83517182.16.A.G.A | cds-KAK7156679.1 | Promoter | CM074871.1 | NA | NA |
| X83517419.41.A.G.A | cds-KAK7124665.1 | Promoter | CM074885.1 | NA | NA |
| X83517765.32.C.A.C | cds-KAK7122878.1 | Promoter | CM074886.1 | NA | NA |
| X83520796.65.T.C.T | cds-KAK7124734.1 | Promoter | CM074885.1 | NA | NA |
| X83522800.10.G.A.G | cds-KAK7144674.1 | Promoter | CM074876.1 | NA | NA |
| X83522942.48.G.A.G | cds-KAK7151378.1 | Promoter | CM074873.1 | P34996.1 | P2Y purinoceptor 1 |
| X83524112.64.C.T.C | cds-KAK7126732.1 | Promoter | CM074884.1 | Q1LXI5.2 | Large ribosomal subunit<br>protein mL54 |
| X83526518.29.G.T.G | cds-KAK7117804.1 | Promoter | CM074890.1 | NA | NA |

**Table S4.**
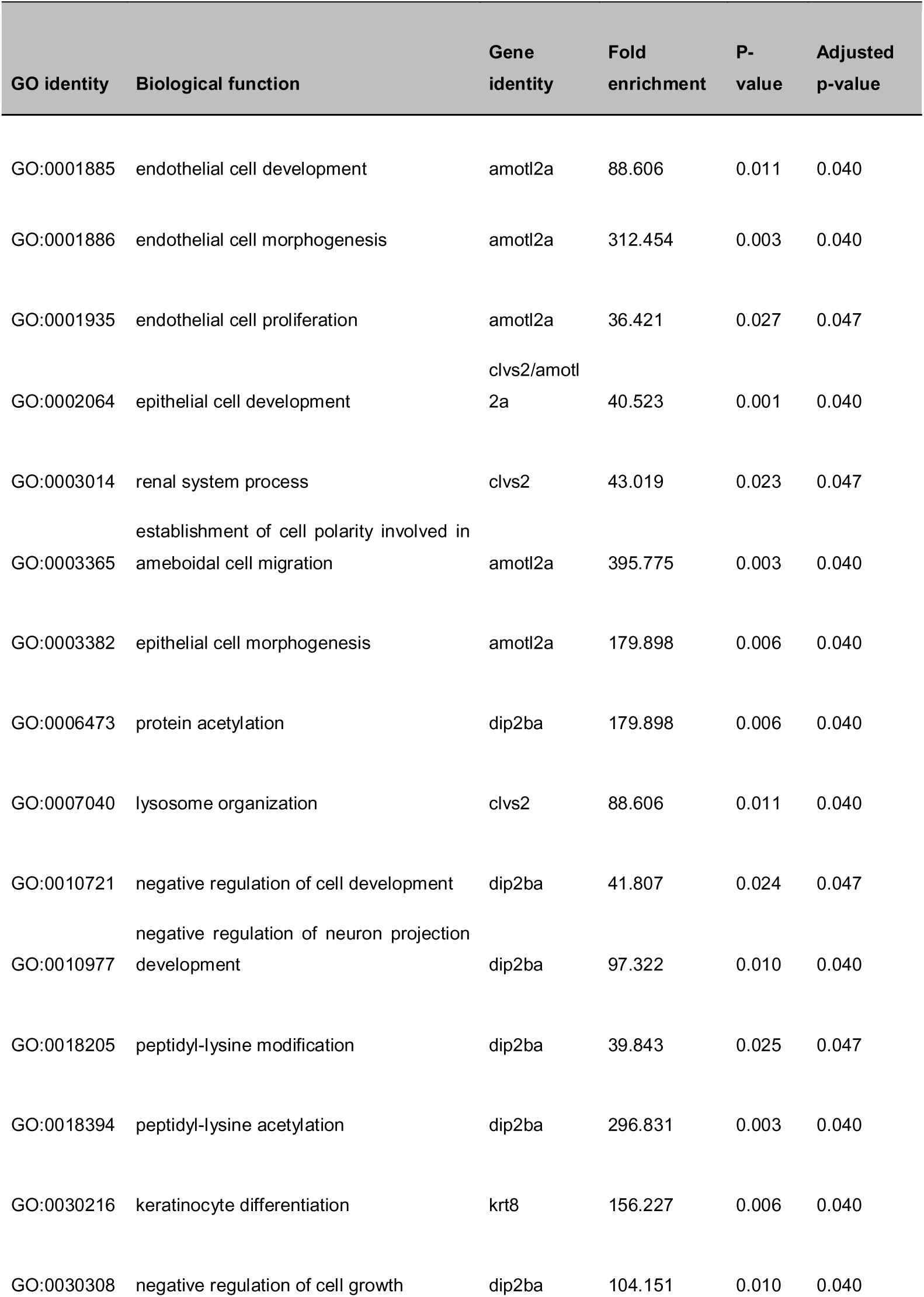
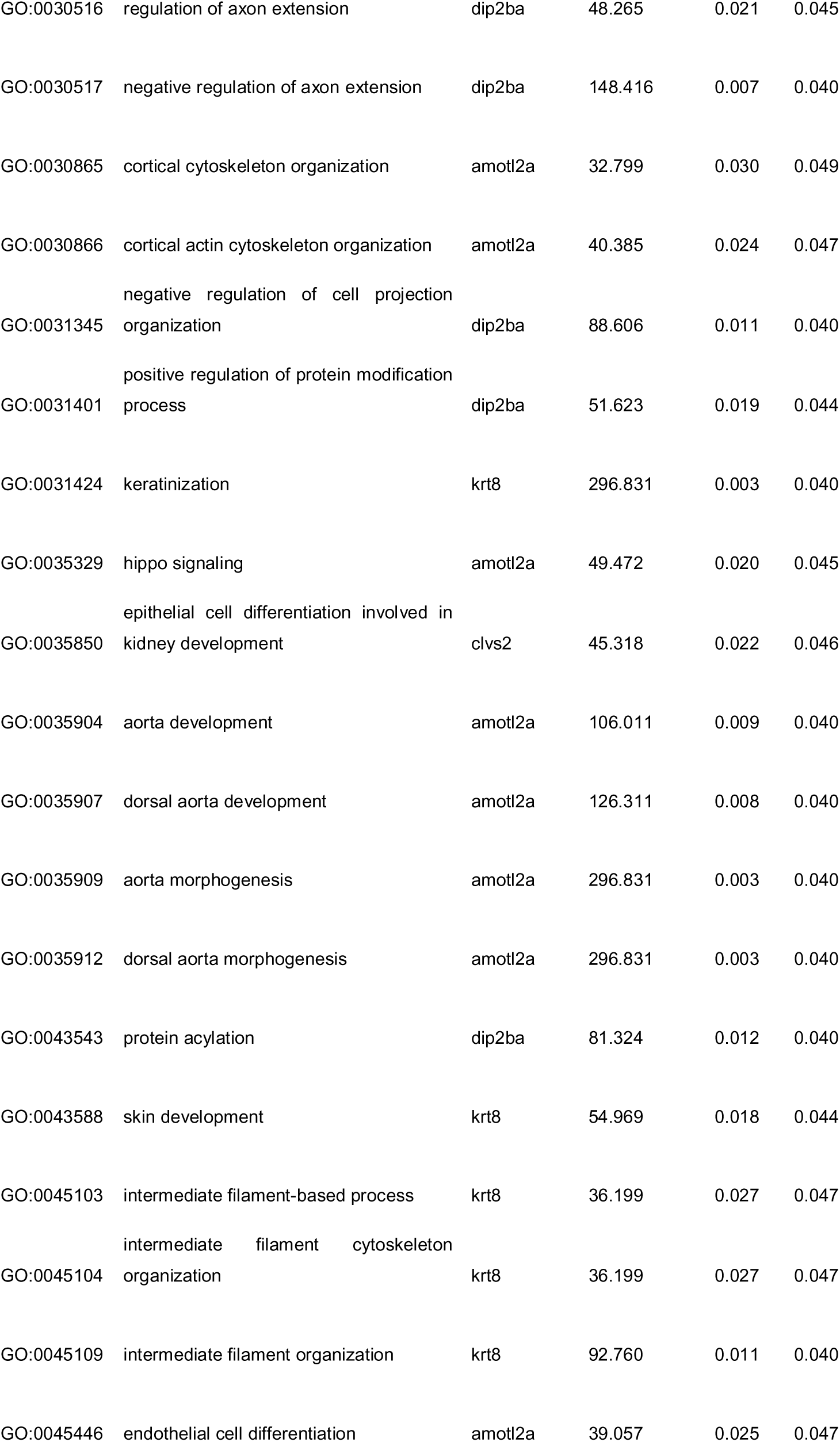
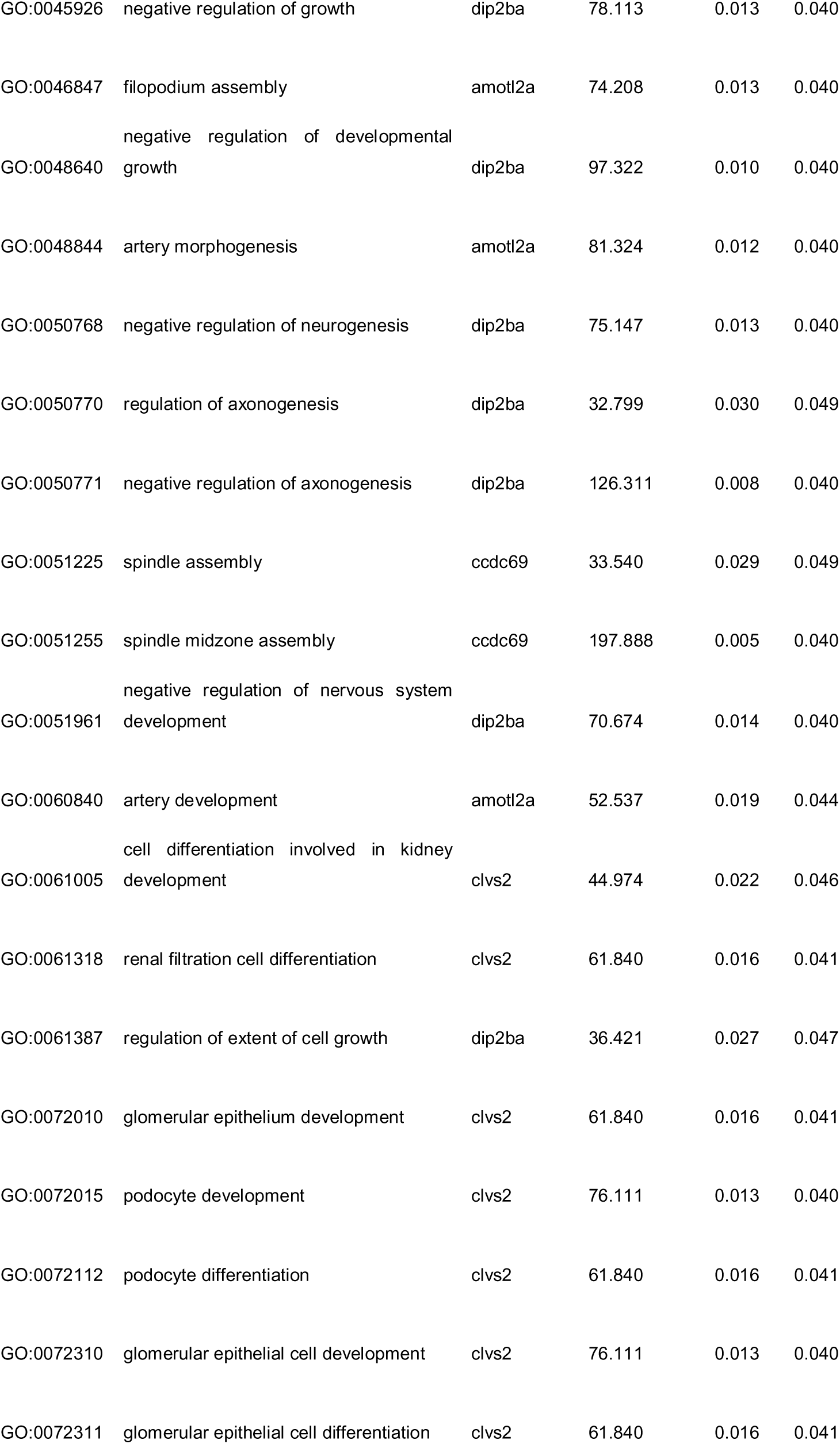
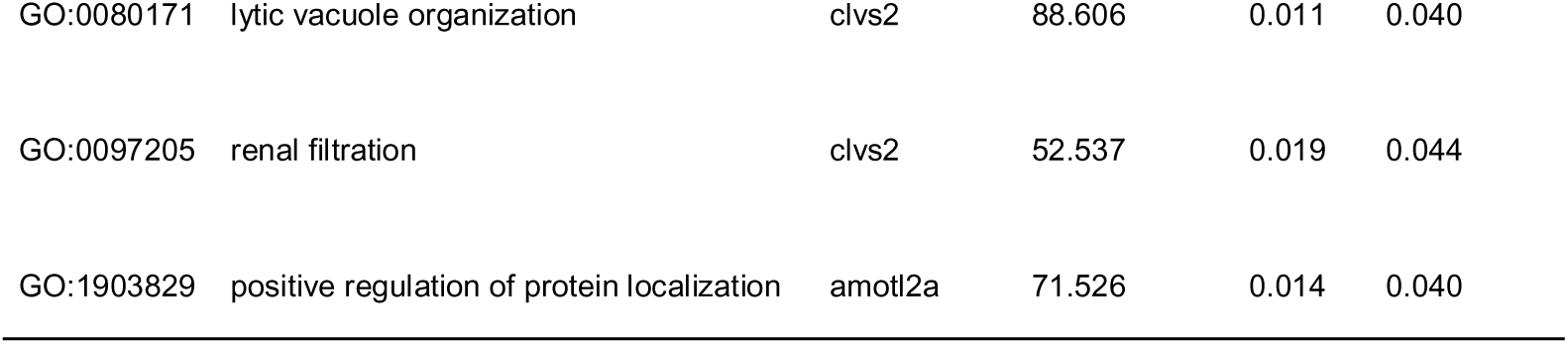
Gene Ontology enrichment analysis of candidate genes associated with ecological responses. Gene Ontology (GO) enrichment analysis was performed on the 21 annotated genes associated with candidate SNPs located within coding sequences (CDS) or promoter regions (Table S3). The table reports significantly enriched Biological Process GO terms, their associated genes, fold enrichment values, raw *P*-values, and Benjamini–Hochberg adjusted *P*-values. Only GO terms with an adjusted *P*-value < 0.05 are reported.

## REFERENCES

1. Hooper, D. U. et al. Effects of biodiversity on ecosystem functioning: A consensus of current knowledge. Ecol. Monogr. 75, 3–35 (2005).

2. Paine, R. T. Food Web Complexity and Species Diversity. Am. Nat. 100, 65–75 (1966).

3. Loreau, M. From Populations to Ecosystems: Theoretical Foundations for a New Ecological Synthesis. (Princeton University Press, 2010).

4. Des Roches, S. et al. The ecological importance of intraspecific variation. *Nat*. Ecol. Evol. 2, 57–64 (2018).

5. Raffard, A., Santoul, F., Cucherousset, J. & Blanchet, S. The community and ecosystem consequences of intraspecific diversity: a meta-analysis. Biol. Rev. 94, 648–661 (2019).

6. Fargeot, L. et al. Genetic diversity affects ecosystem functions across trophic levels as much as species diversity, but in an opposite direction. eLife 13, RP100041 (2025).

7. Harmon, L. J. et al. Evolutionary diversification in stickleback affects ecosystem functioning. Nature 458, 1167–1170 (2009).

8. Farkas, T. E., Mononen, T., Comeault, A. A., Hanski, I. & Nosil, P. Evolution of Camouflage Drives Rapid Ecological Change in an Insect Community. Curr. Biol. 23, 1835–1843 (2013).

9. Crutsinger, G. M. et al. Plant Genotypic Diversity Predicts Community Structure and Governs an Ecosystem Process. Science 313, 966–968 (2006).

10. Prunier, J. G. et al. Genetic erosion reduces biomass temporal stability in wild fish populations. Nat. Commun. 14, 4362 (2023).

11. Leigh, D. M., Hendry, A. P., Vázquez-Domínguez, E. & Friesen, V. L. Estimated six per cent loss of genetic variation in wild populations since the industrial revolution. Evol. Appl. 12, 1505–1512 (2019).

12. Exposito-Alonso, M. et al. Genetic diversity loss in the Anthropocene. Science 377, 1431–1435 (2022).

13. Whitham, T. G. et al. A framework for community and ecosystem genetics: from genes to ecosystems. Nat. Rev. Genet. 7, 510–523 (2006).

14. Hughes, A. R., Inouye, B. D., Johnson, M. T. J., Underwood, N. & Vellend, M. Ecological consequences of genetic diversity: Ecological effects of genetic diversity. Ecol. Lett. 11, 609–623 (2008).

15. Hughes, A. R. & Stachowicz, J. J. Genetic diversity enhances the resistance of a seagrass ecosystem to disturbance. Proc. Natl. Acad. Sci. 101, 8998–9002 (2004).

16. Reusch, T. B. H., Ehlers, A., Hämmerli, A. & Worm, B. Ecosystem recovery after climatic extremes enhanced by genotypic diversity. Proc. Natl. Acad. Sci. 102, 2826–2831 (2005).

17. Barbour, M. A. et al. Genetic specificity of a plant–insect food web: Implications for linking genetic variation to network complexity. Proc. Natl. Acad. Sci. 113, 2128–2133 (2016).

18. Skovmand, L. H. et al. Keystone Genes. Trends Ecol. Evol. 33, 689–700 (2018).

19. Barbour, M. A., Kliebenstein, D. J. & Bascompte, J. A keystone gene underlies the persistence of an experimental food web. Science 376, 70–73 (2022).

20. Blanchet, S., Fargeot, L. & Raffard, A. Phylogenetically-conserved candidate genes unify biodiversity–ecosystem function relationships and eco-evolutionary dynamics across biological scales. Mol. Ecol. 32, 4467–4481 (2023).

21. Schweitzer, J. A. et al. Genetically based trait in a dominant tree affects ecosystem processes. Ecol. Lett. 7, 127–134 (2004).

22. Crutsinger, G. M. et al. Testing a ‘genes-to-ecosystems’ approach to understanding aquatic-terrestrial linkages. Mol. Ecol. 23, 5888–5903 (2014).

23. Wuest, S. E. & Niklaus, P. A. A plant biodiversity effect resolved to a single chromosomal region. *Nat*. Ecol. Evol. 2, 1933–1939 (2018).

24. Wuest, S. E. et al. Single-gene resolution of diversity-driven overyielding in plant genotype mixtures. Nat. Commun. 14, 3379 (2023).

25. Forsman, A. & Wennersten, L. Inter-individual variation promotes ecological success of populations and species: evidence from experimental and comparative studies. Ecography 39, 630–648 (2016).

26. Osorio, V. et al. Non-native minnows cause much larger negative effects than trout on littoral macroinvertebrates of high mountain lakes. Biol. Conserv. 272, 109637 (2022).

27. Raffard, A., Cucherousset, J., Santoul, F., Di Gesu, L. & Blanchet, S. Climate and intraspecific variation in a consumer species drive ecosystem multifunctionality. Oikos 2023, e09286 (2023).

28. Stanislawek, M. et al. Global Warming and Genomic Diversity Loss Alter the Biomass and the Size Distribution of Experimental Fish Populations. Glob. Change Biol. 31, e70611 (2025).

29. Richard, M. et al. The Aquatic Metatron: A large-scale experimental facility to study the combined effects of habitat fragmentation and climate change on aquatic meta-ecosystems. Methods Ecol. Evol. 16, 57–65 (2025).

30. Sansaloni, C. et al. Diversity Arrays Technology (DArT) and next-generation sequencing combined: genome-wide, high throughput, highly informative genotyping for molecular breeding of Eucalyptus. BMC Proc. 5, P54, 1753–6561-5-S7-P54 (2011).

31. Raffard, A. et al. Intraspecific diversity loss in a predator species alters prey community structure and ecosystem functions. PLOS Biol. 19, e3001145 (2021).

32. Oriowo, T. O. et al. A chromosome-level, haplotype-resolved genome assembly and annotation for the Eurasian minnow (Leuciscidae: *Phoxinus phoxinus* ) provide evidence of haplotype diversity. GigaScience 14, giae116 (2025).

33. Loreau, M. Biodiversity and ecosystem functioning: A mechanistic model. Proc. Natl. Acad. Sci. 95, 5632–5636 (1998).

34. Whitham, T. G. et al. Extending Genomics to Natural Communities and Ecosystems. Science 320, 492–495 (2008).

35. Intergovernmental Panel On Climate Change (Ipcc). (2023). Climate Change 2021 – The Physical Science Basis: Working Group I Contribution to the Sixth Assessment Report of the Intergovernmental Panel on Climate Change (1st ed.). Cambridge University Press. 10.1017/9781009157896

36. Schoener, T. W. The Newest Synthesis: Understanding the Interplay of Evolutionary and Ecological Dynamics. Science 331, 426–429 (2011).

37. Boyle, E. A., Li, Y. I. & Pritchard, J. K. An Expanded View of Complex Traits: From Polygenic to Omnigenic. Cell 169, 1177–1186 (2017).

38. Visscher, P. M. et al. 10 Years of GWAS Discovery: Biology, Function, and Translation. Am. J. Hum. Genet. 101, 5–22 (2017).

39. Matthews, B. et al. Toward an integration of evolutionary biology and ecosystem science. Ecol. Lett. 14, 690–701 (2011).

40. Post, D. M. & Palkovacs, E. P. Eco-evolutionary feedbacks in community and ecosystem ecology: interactions between the ecological theatre and the evolutionary play. Philos. Trans. R. Soc. B Biol. Sci. 364, 1629–1640 (2009).

41. He, J., Guo, D., Zhang, L. & Wang, X. Identification of Krt8 as a Newly Discovered Immune-Related Protein in Zebrafish. Preprint at 10.2139/ssrn.4546743 (2023).

42. Kimura, Y. & Nikaido, M. Conserved keratin gene clusters in ancient fish: An evolutionary seed for terrestrial adaptation. Genomics 113, 1120–1128 (2021).

43. Rudman, S. M. et al. What genomic data can reveal about eco-evolutionary dynamics. *Nat*. Ecol. Evol. 2, 9–15 (2018).

